# Structure and Dynamics of a Long-Acting Insulin Analog in Hexameric and Dihexameric States

**DOI:** 10.64898/2026.04.23.720485

**Authors:** Esra Ayan, Hoang Nguyen, Hasan Demirci, Türkan Haliloğlu, Ivet Bahar

## Abstract

Elucidating the structure and dynamics of insulin and its analogs has been of broad interest, while presenting challenges due to the unique structural dynamics of insulin (composed itself of two multiply cross-linked peptides A and B) and its ability to assemble in a variety of oligomeric structures under physiological conditions. Here, we present two distinct X-ray crystallographic structures of the long-acting human insulin analog detemir (INSD) resolved in hexameric and dodecameric (or dihexameric) states at 2.85 Å and 2.70 Å resolution, respectively, using diffraction data collected under ambient temperature conditions. Characterization of the collective dynamics of these oligomers using the Gaussian Network Model (GNM) reveals several key features: (i) Oligomerization imparts high cooperativity in structural dynamics evidenced by dissection of the cross-correlations at various hierarchical levels; (ii) detemir monomers’ conformational flexibility is highly suppressed within oligomeric constructs, the effect being particularly strong in the dihexamer due to the asymmetric packing of the hexamers and the presence of myristoyl groups at B peptides termini whose interactions imparts further heterogeneities; and (iii) a number of key residues retain, however, their intrinsic dynamics, to be deployed upon release from the oligomers. We distinguish in particular residues serving as hinge sites that mediates the conformational dynamics of the asymmetric units (dimers) and monomers (I2_A_–V3_A_ and Y19 _A_ –C20_A,_ and L11_B_-L15_B_ and Y26_B_ of the respective peptides A and B), or as anchors supporting structural stability (disulfide-bridge forming cysteines, plus selected residues such as L16_A,_ G8_B_ and R22_B_-F24_B_. Overall, this study provides a structural–dynamic framework for gaining new insights into the dynamics of long-acting analog INSD and helps identify actionable sites for modulating insulin (analogs) dynamics toward designing more effective therapeutics.

## Introduction

Human insulin is predominantly found in equilibrium between two distinct states, monomeric (5.8 kDa) and dimeric (11.6 kDa), in mildly acidic aqueous media [1]. Each monomer consists of two peptides, chains A and B (of 21 and 30 amino acids, respectively), cross-linked by disulfide bridges. In the presence of ligands such as Zn^2+^ and Cl^-^ ions and phenol, insulin preferably forms a hexameric structure (of 36 kDa) and can assume a variety of oligomeric states under physiological conditions [2]. The degree of self-assembly or multimerization is a key determinant of insulin’s pharmacological profile and clinical efficacy [3]. Favoring the monomeric state enables the design of insulin analogs with rapid therapeutic action, whereas hexameric and higher-order oligomeric states usually induce long-acting bioactivity [3, 4].

X-ray crystallographic studies on hexameric assemblies of insulin go back to the original work of Adams et al in 1969 [5], and many structures of hexameric insulin and its analogs have been resolved since then [4]. Canonical human/bovine insulin hexamers include the 4-Zn^2+^ (T_3_R_3_-like) form [6, 7] as well as the cresol-bound R_6_ form [8]; modified insulins also crystallize as hexamers and dihexamers [9, 10]. Especially, acylated long-acting analogs (e.g., detemir, and more recently, degludec) further exhibit ligand-controlled (mainly zinc and phenol) *dihexameric* assemblies, i.e., dodecamers composed of two hexameric substructures [11, 12].

The engineering of long-acting insulin analogs has been facilitated by advances in genetic engineering and recombinant DNA technology. These advances have led to their approval for type 1 and type 2 diabetes mellitus therapy, beginning with glargine in 2000, shortly followed by detemir (brand name Levemir™, produced by Novo Nordisk until Dec 2024) launched in 2005. These analogs have been shown to provide a steady supply of insulin for extended durations. Insulin detemir (hereafter INSD), indicated for glycemic control [13], proved to control blood sugar levels over a 24-hour period. Its prolonged action has been attributed to its propensity to form a dihexameric form, endowed by covalent bonding of a myristoyl fatty acid chain (of 14 carbon atoms) to residue K29 in peptide B after removal of the C-terminal residue T30. The presence of such an aliphatic fatty acid appears to enhance hydrophobicity in favor of oligomerization. This modification further enables INSD to bind to albumin, but only to a small fraction of serum at therapeutic concentrations, reducing the risk of interference with other drugs that bind to albumin [14]. Comparative kinetic studies and euglycemic clamp analyses have confirmed the prolonged action of INSD, as well as its superior pharmacokinetics and temporal profiles [15]. While prolonged action of INSD has been attributed to oligomerization and reversible albumin binding, the structural properties and intrinsic dynamics of the oligomers, which may support its long-acting effect have been elusive.

Here, we present two X-ray crystallographic structures of INSD, one hexameric, the other dihexameric. In contrast to the previous resolutions of the dihexameric insulins or analogs under cryogenic temperatures and from single crystals [9, 12], the current structures have been resolved at ambient temperature (293K) and are derived from multiple crystals. Furthermore, the present hexameric structure of detemir is the first ‘isolated hexamer’ observed by X-ray diffraction to stably form under ambient temperature; whereas previously determined hexamers have been separated from the cryogenic dihexamer [16].

The new structural data, which more closely represent physiological conditions, are used to perform a comparative analysis of the intrinsic dynamics of detemir across multiple scales, from monomer, to dimer, hexamer, and dihexamer, using the Gaussian Network Model (GNM) [17–20]. GNM yields a unique analytical solution for the spectrum of motions (normal modes) accessible to proteins, based on their specific inter-residue contact topology. Cross-correlations predicted by the GNM between the equilibrium fluctuations of structural elements at multiple scales often explain the molecular basis of allosteric events in multimeric systems [21], as well as the role of oligomerization in defining biological function [22, 23].

The current GNM analysis of INSD provides a structure-based understanding of the equilibrium dynamics of detemir oligomers and sheds light on the effect of oligomerization on the structural dynamics of the monomers. It also points to the intrinsic characteristics of the insulin monomer that are maintained in the oligomeric constructs, upon the dissociation into dimers and monomers, long-acting action of this engineered insulin. Our study highlights the identity of key residues (e.g., global hinge sites and local energy concentration centers) that underlie the dynamics and stability of insulin or detemir monomers and dimers, which can potentially serve as targets for modulating the function of insulin or engineering novel insulin analogs.

## Results

### Resolution of detemir dihexamer and hexamer near-physiological structures

We determined the near-physiological structure of dihexameric INSD at 2.7 Å resolution, and that of hexameric INSD at 2.88 Å. The diffraction data were collected from 72-well Terasaki microbatch screening plates at the Turkish Light Source by directly mounting the plates onto the *XtalCheck* plate reader adapter module [24, 25]. **Fig. 1** displays the resolved structures and electron density maps for INSD in hexameric and dihexameric states, and **Table 1** lists the corresponding properties and refinement statistics.

**Figure 1.**
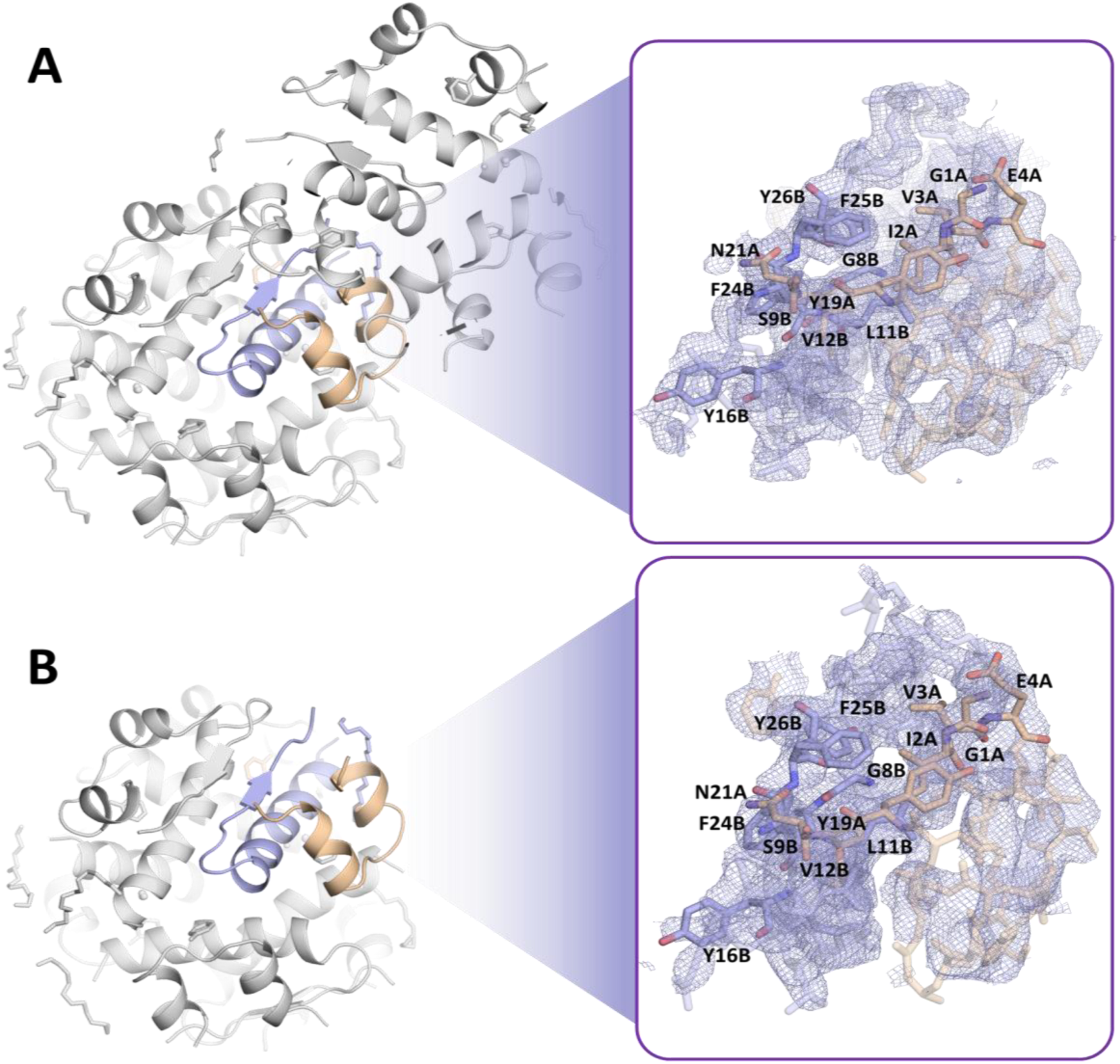
Dihexameric and hexameric structures resolved for INSD. The structures are shown in cartoon (50% transparency) representation, and the chains A and B of an INSD insulin monomer are shown in *wheat* and *light blue*, respectively. Representative 2Fo-Fc electron density maps contours at the 1σ level are displayed in the right panels to illustrate the overall quality and interpretability of the model. Insulin residues observed in prior cryo-EM studies to interact with the insulin receptor (IR) are labeled in the insets to provide structural context within the present model.

**Table 1.** Data collection and refinement statistics.

| Data collection | Detemir-hexamer | Detemir-dihexamer |
| --- | --- | --- |
| PDB ID | 9LVY | 9LVX |
| Instrument | Turkish DeLight | Turkish DeLight |
| Resolution range (Å) | 17.56 - 2.85 (2.95 - 2.85) | 18.89 - 2.70 (2.80- 2.70) |
| Space group | R 3: H | R 3: H |
| Unit cell | 81.239 81.239 40.321 90 90 120 | 80.898 80.898 80.414 90 90 120 |
| Total reflections | 48296 (3457) | 102108 (8477) |
| Unique reflections | 2306 (202) | 5361 (479) |
| Redundancy | 19.0 (16.5) | 12.9 (11.1) |
| Completeness (%) | 98.53 (94.01) | 97.30 (93.40) |
| Mean I/sigma(I) | 7.39 (2.39) | 5.81 (2.57) |
| Wilson B-factor | 53.57 | 34.78 |
| CC1/2 | 0.89 (0.745) | 0.884 (0.737) |
| CC* | 0.971 (0.716) | 0.969 (0.921) |
| R-work | 0.2854 (0.3990) | 0.2734 (0.3511) |
| R-free | 0.3369 (0.4509) | 0.3259 (0.3337) |
| No. of reflections in ref. | 31770 (2244) | 69174 (5706) |
| Protein residues | 100 | 200 |
| RMS (bonds) | 0.002 | 0.002 |
| RMS (angles) | 0.44 | 0.40 |
| Ramachandran favored (%) | 100.00 | 99.46 |
| Ramachandran allowed (%) | 0.00 | 0.54 |
| Ramachandran outliers (%) | 0.00 | 0.00 |
| Average B-factor | 65.67 | 46.98 |
| macromolecules | 63.15 | 41.49 |
| ligands | 108.43 | 55.12 |
| solvent | 39.90 | 99.60 |

The INSD hexamer exhibited distinct three-dimensional orientations within the asymmetric unit cell, adopting an isolated “dimeric” arrangement with unit cell dimensions (ASU) specified as a = b = 80 Å, and c = 40 Å and α = β = 90° and γ =120° (**Fig. 1B**); whereas the dihexameric form adopted an isolated dimer-of-dimer assembly in the asymmetric unit, characterized by unit cell dimensions of a = b = c = 80 Å, with α = β = 90° and γ =120° (**Fig. 1A**). Both conformers were classified within the R3:H space group. The dihexamer structure (in ASU) contained four myristic acid moieties, four phenol molecules, four Zn^2+^ cations, and four Cl^-^ anions, whereas the hexamer form (in ASU) had two of each.

The insets in **Fig. 1** indicate residues observed in previous studies [26–29] to engage the insulin receptor (IR). Minor rearrangements are discerned in the side chains of these residues in the hexameric and dihexameric forms. Two binding sites have been observed in previous cryo-EM structures [29]: a high-affinity site (Site 1) that includes G1_A_, I2_A_, V3_A_, E4_A_, Y19_A_, and N21_A_ on chain A, and V12_B_, Y16_B_, G23_B_, F24_B_, F25_B_, and Y26_B_ on chain B; and a low-affinity site (Site 2) comprising S12_A_, L13_A_, and E17_A_ (chain A) and H10_B_, E13_B_, and L17_B_ (chain B) [29, 30]. The conformational dynamics of these residues in the oligomers, as well as in the dimeric state as an intermediate before insulin function, and in the monomeric (functional) form, will be characterized by the GNM.

The structures of the hexamers in the hexameric and dihexameric states revealed a striking degree of similarity within their asymmetric units (root-mean-square-deviation (RMSD) of 0.384 Å) (**Fig. S1A**). Furthermore, the individual monomers’ backbones were closely superimposable (**Fig. S1B**). In both cases, the hexamers were composed of triplets of dimers. We note that all three dimers are structurally identical in the hexamer. However, in the dihexamer, the front-to-back association of the hexamers (**Fig. 2A**) entails asymmetric inter-hexamer contacts and myristoylation groups interactions, which result in two types of dimers subject to different quaternary interactions, which will be referred to as Type 1 and Type 2 dimers.

**Figure 2.**
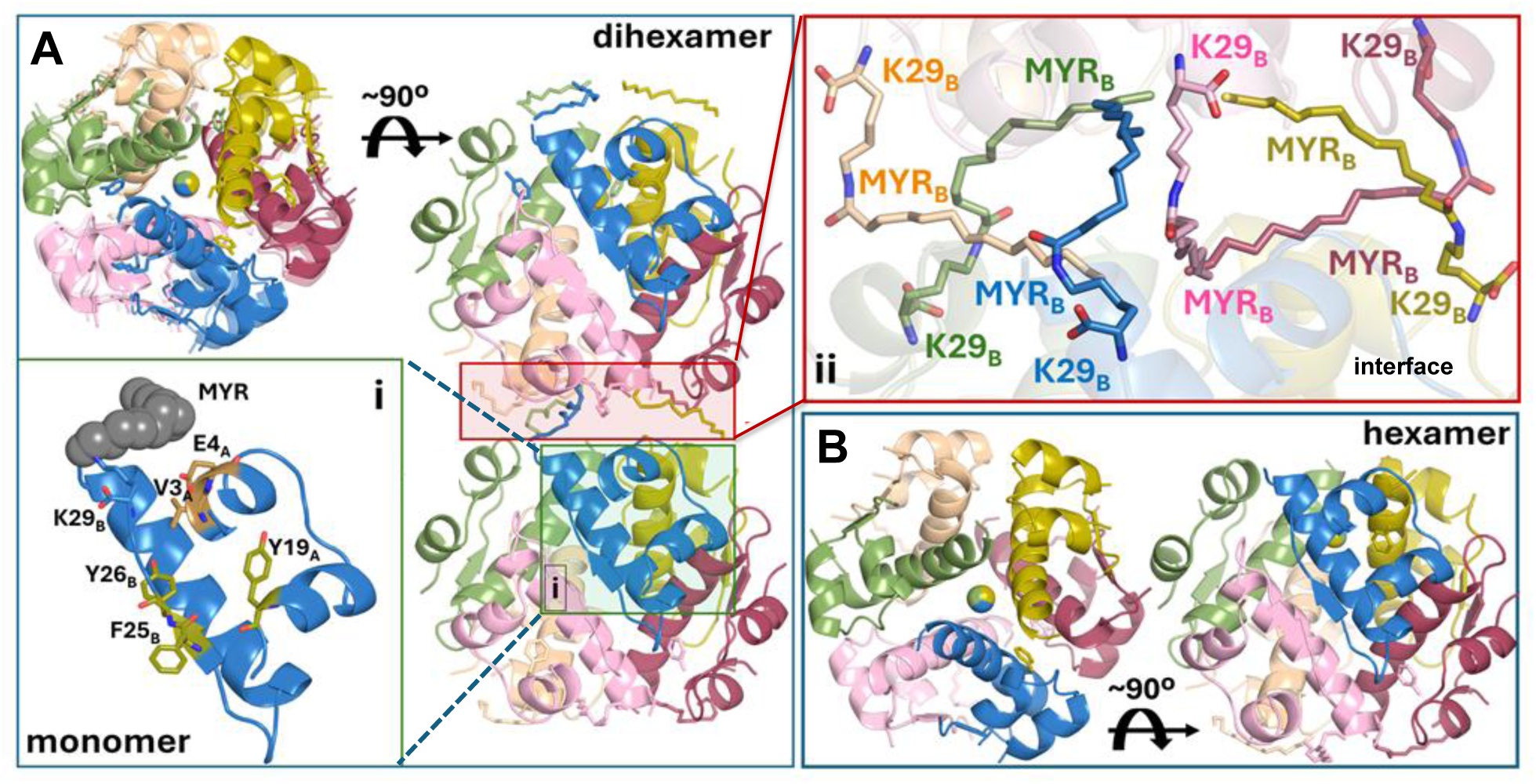
Packing of monomers in the dihexameric and hexameric INSDs, and the role of myristoyl chains at dihexamer interface. **(A)** Two orthogonal views of the dihexamer (∼90° rotation) reveal the packing geometry of the hexamers against each other at their myristoylated ends (enclosed in *pink box*). Two insets are shown, (i) on the *lower left,* and (ii) on the *upper right*. The inset (i) provides a closeup view of a single monomer with the myristoyl (MYR) moiety (*gray space-filling*) covalently linked to K29_B_. A few residues (E4_A_, V3_A_, Y19_A_, F25_B_, Y26_B_) known to interact with the receptor are labeled (*see also,* Figure 1). The inset (ii) displays a closeup view of the inter-hexamer interface, illustrating how the MYR groups of adjacent hexamers cluster together to stabilize the dihexamer. **(B)** For comparison, two orthogonal views of the isolated hexamer are shown using the same color code as the hexameric parts of the dihexamers (∼90° rotation).

**Fig. 2A** shows the dihexamer structure from two perspectives (*top* and *side views*). An enlarged view of one of the monomers (*blue ribbon*, inset (i) on the *lower left*) illustrates the proximity of the myristoyl group to V3_A_ and E4_A_. The aromatic side chains (Y19_A_, F25_B_, and Y26_B_) which participate in IR binding make intermonomer contacts (across antiparallel β-strands belonging to adjacent monomers) in the dimers that assemble into the hexamer. The second inset (*upper right*, inset ii) shows a close-up view of the interface between the two hexameric subunits of the dihexamer. A tight web of interactions between six myristoyl chains, three contributed by each hexamer, glues the hexameric subunits together. Comparison of **Figs. 2A** and **2B** further illustrates the structural similarity between the hexamers resolved in the isolated state (**Fig. 2B**) and as part of a dihexamer (**Fig. 2A**). As will be shown below, participation in oligomers of different sizes significantly alters the exposure of critical residues, as well as the dynamics of individual monomers even as the substructures retain similar structural characteristics.

### Dihexameric assembly exhibits a significantly higher cooperativity than that observed in hexameric state

The dihexameric assembly is known to be significantly more effective than the hexameric form in conferring the long-acting activity of insulin. Toward gaining a mechanistic understanding of the basis for the distinct functional behaviors of the INSD hexamer and dihexamer, we analyzed their intrinsic dynamics using the GNM. GNM provides information on the equilibrium fluctuations of residues as well as their cross-correlations under physiological conditions. One can dissect the accessible spectrum of collective motions to distinguish the ‘global motions’ (driven by the low frequency modes) as well as the highest-frequency fluctuations (that point to centers of energy localization) [17–20]. The low frequency modes (also called *soft modes*) are particularly interesting as they are the most readily accessible motions (requiring the lowest energy ascent for a given structural change) that cooperatively engage large substructures essential to enabling functional events [31, 32] or to eliciting allosteric responses [33]. Residues distinguished by their high mobility in the high-frequency modes, on the other hand, are often termed “kinetically hot residues”, and they are subject to high frequency motions due to their tight packing; hot residues often form folding nuclei or regulate conserved interactions [34–36]. Previous GNM studies have demonstrated how a protein’s dynamics within a complex or biological assembly differ from that in isolation. Participation in a restrictive environment affects the protein dynamics by selectively inhibiting or modulating certain types of motion [37]. Examination of the dynamics of intact structures, rather than substructures, is therefore paramount to making inferences on functional mechanisms. This is particularly important, as multimerization may favor or impede signaling events and cellular responses [22]. Previous work has also demonstrated that despite the alterations caused by complexation or multimerization, the building blocks still retain some intrinsic features, which may even be amplified in symmetric assemblies [33] or in tandem repeats [38]. In this respect, it is useful to examine the intrinsic dynamics of the building blocks (e.g., monomers or dimers) in the context of the assembly toward elucidating the conserved features which are sustained in the oligomers and would become operative upon disassembly.

Based on these considerations, we analyzed the conformational dynamics of INSD molecules under various multimerization states (*colored pale blue* in **Fig. 3A**), ranging from di-hexamer (*leftmost*) to isolated monomer (*rightmost*). We evaluated in each case the structural dynamics of the multimers as well as that of an embedded monomer (*shown in dark blue in the ribbon diagrams*). The heat maps in **Fig. 3B-C** describe the cross-correlations between the motions of all pairs of residues, as defined by the global dynamics of each multimer (**B**) and to a monomer embedded in that multimer (**C**). Global dynamics refers to movements driven by the softest collective modes. To be consistent across different multimers, we selected a subset of soft modes that ensures a cumulative variance of ≥0.40 in all the four cases (dihexamer, hexamer, dimer, and monomer) (see Methods and **Fig. S2**). The heatmaps are color-coded by the correlation cosines C*_ij_* between the motions of residues *i* and *j*, in these global motions. *C_ij_* varies in the range [−1, 1], with the lower and upper limits corresponding to the fully anticorrelated (*blue*; coupled but opposite-direction) and fully correlated (*red*; same-direction) movements, respectively; and C*_ij_* = 0 (*green*) refers to uncorrelated pairs (see the *color bar* in **Fig. 3B-C**).

**Figure 3.**
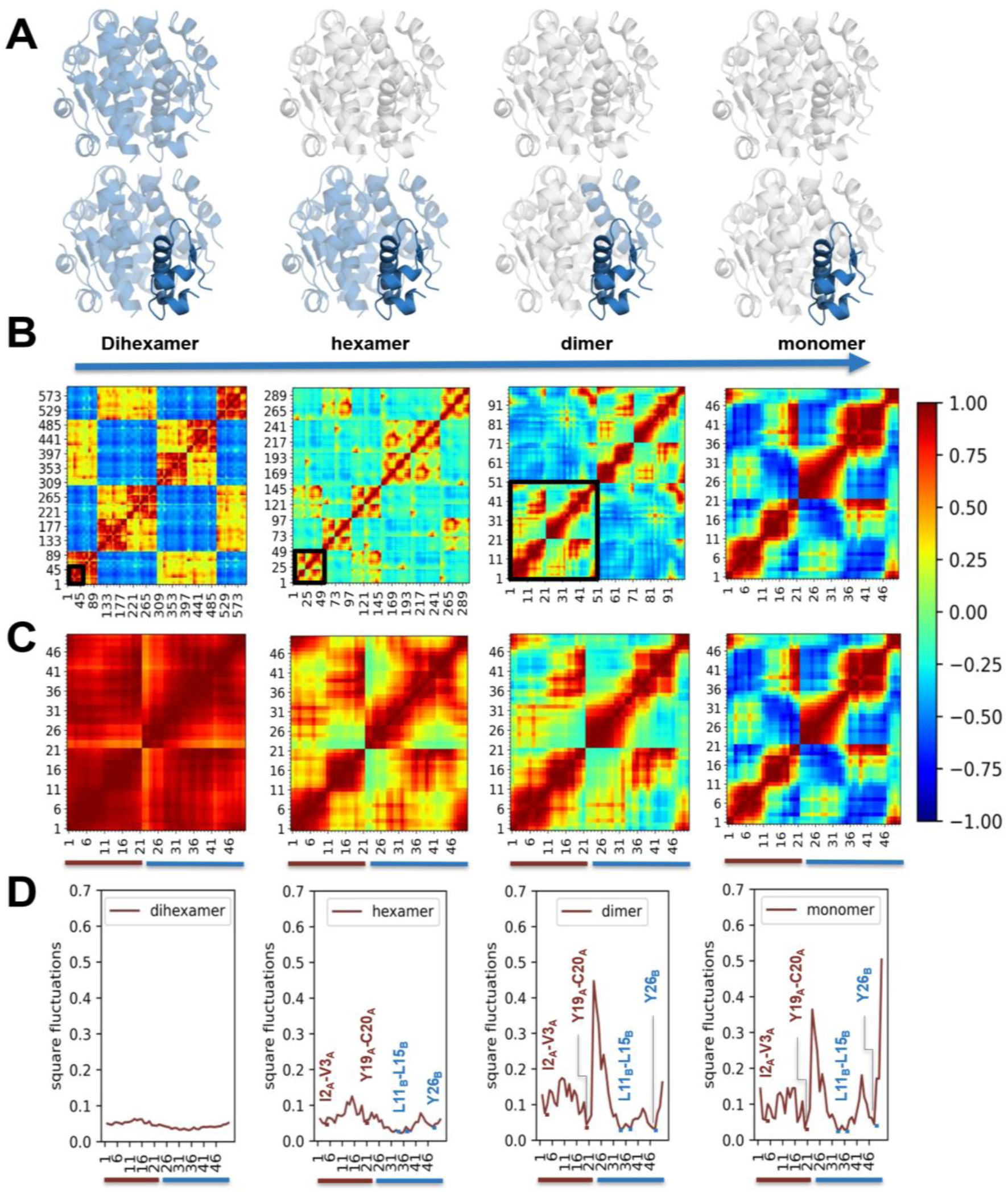
Global dynamics of INSD oligomers and embedded monomers. **(A)** Four different multimerization states of INSD (*in light blue*). **(B)** Cross-correlations between residue motions driven by GNM soft modes in each multimerization state (see the *color code* on the right). The *black square* in each map indicates a monomer, whose cross-correlation behavior is magnified in panel (**C**). The axes refer to residue numbers, mainly residues 1-21 of chain A, and 1-29 of chain B renumbered as 22-50 here and in the following figures. **(D)** Mobility profiles of embedded monomers. The y-axis shows the square fluctuations of monomer residues based on the normalized distribution of global motions accessible to the four cases. Hinges are indicated by *brown* (chain A) and *blue* (chain B) *dots* and *labels*. Hinge regions are observed at the two terminal short helices of chain A (I2_A_–V3_A_ and Y19_A_ –C20_A_), at the central portion of the long helix (residues L11_B_–L15_B_), and at the spatially neighboring Y26_B_ (in the β-strand at the C-terminal segment) in chain B. See the location of these residues in the detemir monomer in Fig. 4.

The cross-correlation maps in **Fig. 3B** reveal the regions that concertedly move in the same direction *(red* blocks along the diagonal), as well as those structural elements that undergo strongly anticorrelated movements (*blue blocks*), or those weakly correlated (*yellow*) or uncorrelated (*green*). Comparison of the maps for the dihexamer (1 ≤ *i* ≤ 600; *leftmost* in panel **B**) and hexamer (1 ≤ *i* ≤ 300; *2^nd^ map* from *left*) shows that the motions within the dihexamer are significantly more coherent and more cooperative than those within the hexamer. This is evidenced by sharp *red* and *blue* regions in the dihexamer (indicative of strong couplings, positive or negative, respectively) as opposed to the dominance of weakly correlated (*yellow* regions) uncorrelated movements (*green* regions) within the hexamer. Thus, *oligomerization into a dihexamer confers significant cooperativity across the monomers (and dimers),* as compared to oligomerization into a hexamer.

Closer examination of **Fig. 3B** shows couplings *within* the individual monomers along the diagonal of the hexamer heat map; however, the couplings *across* monomers are weak to moderate. This contrasts with the behavior of the dihexamer, which cooperatively engages multiple monomers. Notably, in the dihexamer, the dimers composed of monomers (1,2), (3,4), and (5,6) within each hexameric subunit show high intra-dimer correlations, and strong inter-dimer anticorrelations including those between different hexameric subunits. Such cooperative movements across dimers are hardly observable in the soft modes of the hexamer.

As a further test, we considered tetramers composed of pairs of dimers belonging to two hexamers, also called a di-dimer. As shown in the **Fig. S3**, the soft modes in this case closely agree with the portion of the di-hexamer map corresponding to the relative movements of these four monomers, which means that the relative movements of the two dimers dominate the observed motions. This tetrameric state will not be considered further, as it represents an oligomeric state not observed in experiments.

### The intrinsic dynamics of monomeric units are modulated by selected residues that support the insulin conformational mechanics and fold stability

We next focused on the behaviors of monomers embedded within the hexameric and dihexameric assemblies (**Fig. 3C**). For simplicity, we first focus on the first monomer of each structure, enclosed in the *black squares* in **Fig. 3B**. The hexameric units are generated in the R3:H space group by the crystallographic three-fold symmetry applied to a single asymmetric unit containing one dimer; and the monomer analyzed here represents the first monomer of that asymmetric unit. The second monomer, while sharing the exact same structure as the first, experiences different quaternary contacts, resulting in slightly altered dynamics (*see below*). In the case of the dihexamer, the occurrence of asymmetric dimer-dimer contacts (because of di-dimer in ASU) introduces another level of complexity, resulting in two different types of dimers as mentioned above.

The monomer embedded in the dihexamer moves almost as a rigid block (all *red*). Its residue fluctuation profile (**Fig. 3D**, *leftmost panel*) is almost flat, consistent with the *en-bloc* movement of the monomer. This behavior is a consequence of the dominance of the slowest mode in the di-hexamer (contributing alone to 20% of the total variance; See **Fig. S2**). In this mode, the two hexamers undergo rigid-body anticorrelated movements with respect to each other.

The same monomer in the hexamer, on the other hand, exhibits some internal flexibility during the global modes of the hexamer, evidenced by minima and maxima in its mobility profile (**Fig. 3D**, *2^nd^ map from left)*. We distinguish two minima regions in each peptide, I2_A_–V3_A_ and Y19_A_–C20_A_ in peptide A, and L11_B_–L15_B_ and Y26_B_ in peptide B. These are the most severely constrained regions of the monomers in the hexameric environment.

Notably, these minima and the overall fluctuation profile are consistent with those of the isolated monomer (**Fig. 3D**, *rightmost* panel), indicating that the intrinsic dynamics of the isolated monomer is retained, albeit somewhat suppressed, in the soft motions of the hexamer. This general decrease in the amplitudes of motions is a natural consequence of the constrained hexameric environment. We also note the suppression of the movements of chain termini, especially those of peptide B, in the hexameric environment, consistent with the role of myristoylated ends in the hexamer.

Notably, the patterns characteristic of the global mobility profile of the isolated monomer are clearly retained in the dimeric state’s global dynamics (*3^rd^ heat map* in **Fig. 3C** and corresponding fluctuation profile in **Fig. 3D**). The isolated monomer, devoid of any constraints imposed by neighboring monomers, exhibits multiple minima and peaks, in addition to those retained in the dimer and hexamer. A closer examination of the dynamics of the insulin monomer, presented in **Fig. 4**, reveals other minima (C7_A_, C11_A_, and L16_A_) in the global modes (panel **B**), which persist in the spectrum of all modes (panel **A**). Note that the former two are involved in disulfide cross-links C6_A_-C11_A_ and C7_A_-C7_B_ and furthermore, the hinge residue C20_A_ in peptide A also forms a cross-link C20_A_-C19_B_. Finally, the examination of the other end of the spectrum, the highest frequency regime, or fast modes, yields peaks (as the most constrained regions, also called centers of energy localization) at V3_A_, C6_A_, C7_A_, L11_A_, L16_A_ and C20_A_ in peptide A, and G8_B_, L15_B_, C19_B_, and Y26_B_ in peptide B (**Fig. 4C**). Notably, these residues include (i) the hinges identified in slow modes (V3_A_ and C20_A_ in peptide A, and L15_B_ and Y26_B_ in peptide B) further underscoring their importance as anchors mediating the global modes, and (ii) the disulfide-bridge-forming residue pairs C6_A_-C11_A_, C7_A_-C7_B_ and C20_A_-C19_B_; and (iii) an additional residue, G8_B_, in peptide B.

**Figure 4.**
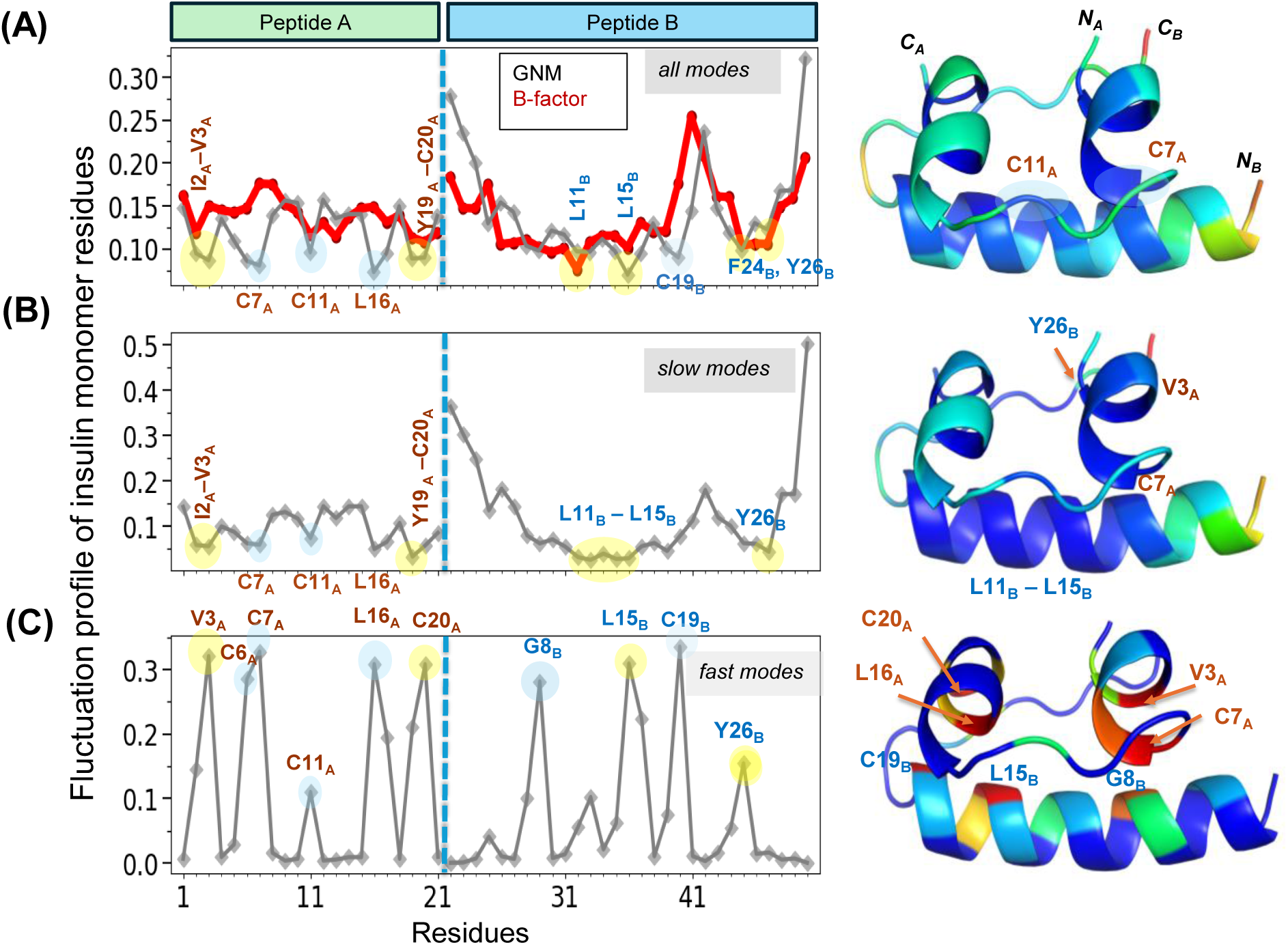
Collective dynamics of insulin monomer. Residue mean-square fluctuation profiles (*gray curves* with *diamond symbols*) predicted by the GNM are shown for both peptides A and B (see *top bar*) based on **(A)** all modes, **(B)** soft/slow modes with a cumulative covariance of >0.40, and **(C)** high frequency modes (fastest 10 modes). Computations were performed using the coordinates extracted from the monomer found in our resolved dihexameric structure, PDB ID: 9LVX. Panel **A** displays the B-factors profile (*red curve*) reported in the PDB. Hinge residues observed in the global modes (panel **B**) are highlighted by *yellow dots*; other highly constrained residues (minima in global modes or maxima in fast modes) are highlighted by *blue dots.* Residues belonging to peptides A and B are labeled in *dark red* and *blue,* respectively. The ribbon diagrams on the *right* visualize regions of high mobility (*red/yellow*) and rigidity (*blue*), consistent with the fluctuation profiles on the *left*. See Fig. 5 for a closer view of the network of interactions between global hinge sites.

In general, minima observed in the GNM global mobility profiles refer to spatially clustered residues that serve as hinges or anchors and cooperatively mediate functional motions. The residues I2_A_–V3_A_, Y19_A_–C20_A_, L11_B_–L15_B_, and Y26_B_ persistently serve this role in the monomer, dimer, and hexamer, and as will be shown in the next subsection, they are hidden but maintained in the dihexamer as well. They are simply not visible in the global modes ‘window’ of the dihexamer. **Fig. 5** illustrates the network of intra-chain and interchain interactions in which these residues are involved. Our analysis consistently suggests that these five residues play an essential role in maintaining/mediating the most cooperative (softest) modes of motion intrinsically accessible to the detemir monomer.

**Figure 5.**
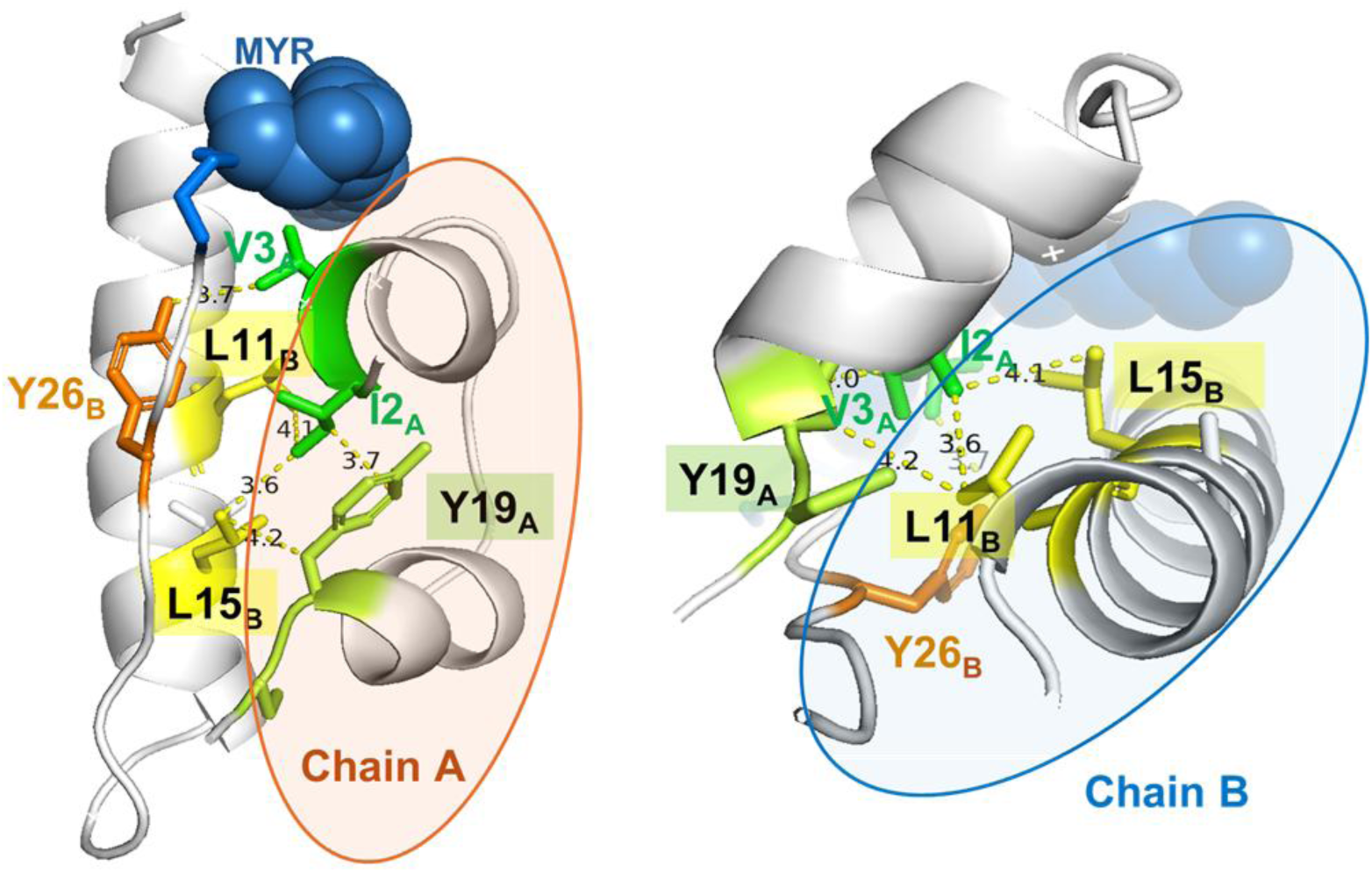
Hinge residues critical to the conformational mechanics of INSD monomer, maintained in oligomeric assemblies. I2_A_ and Y19_A_ make close (< 3.9 Å) intrachain hydrophobic contacts involving three pairs of carbon atoms; L11_B_ and L15_B_ sidechains both interact with I2_A_ sidechain; L15_B_ further interacts with Y19_A_ C_β_; and the hydrophobic core formed by these contacts is consolidated by the close (3.7 Å) interchain association of Y26_B_ with V3_A_.

Obviously, the structural integrity of insulin is ensured by three disulfide bridges (one intra-chain, C6_A_–C11_A_, and two inter-chains, C7_A_–C7_B_ and C20_A_–C19_B_), and that of INSD is further consolidated by the interactions of the C-terminal myristoyl group of peptide B with the N-terminal residue V3_A_ of peptide A. The above analysis shows that at least one of the cysteines in each disulfide bridge also serves as an anchor in the global motions of the monomer (C7_A_, C11_A_, C20_A_). It is worth noting that the hinges and disulfide bridges play different roles: the former group of residues supports the conformational mechanics; and the latter, the fold stability. C20_A_ apparently plays a dual role, as a hinge site and a disulfide bridge-forming residue.

Collectively, the participation in larger oligomeric structures increasingly endows a higher cooperativity and more coherent movements within the monomers; whereas in the dimeric and monomeric forms, the insulin dynamics is expressed with minimal or no constraints. The range of cooperativity is limited to relatively near neighbors along the sequence, as evidenced by the narrower diagonal (*red*) band in their cross-correlations maps (*leftmost maps* in **Fig. 3C**). This observation suggests that the dissociation of the oligomers into dimeric (intermediate) or monomeric (active) forms of the insulin might be entropically more favorable for the dihexamer, compared to the hexamer, due to a higher entropic gain; but may be resisted by an equally high enthalpic cost that stably binds the dimers and monomers. Therefore, the multimers, and in particular the dihexameric state, provide a highly stable environment for packaging the insulin or detemir monomers and retaining the intrinsic dynamics features of key residues to enable the observed long-acting effects while allowing for an entropically-favored release.

### The intrinsic dynamics of the detemir monomers can be discerned in the moderate-to-higher frequency modes of the oligomers

The observed suppression of the monomer mobility in the global dynamics of the dihexamer mainly originates from the prevalence of a new regime of collective modes that operates in this oligomeric state. In this new regime, the soft modes engage large blocks of structures that move in concert, including the entire hexameric subunits, where the individual monomers and dimers move almost rigidly within the hexamers, such that local (intra-monomer) structural changes are hardly detectable. However, upon widening the window of observation to the moderate-to-higher frequency regime of the mode spectrum, the character of the monomers, reminiscent of that of the isolated monomer, emerge, as explained next.

In order to assess whether the intrinsic character of the monomers, not visible in the dihexamer (**Fig. 3C–D**; *left panels*), is still retained albeit hidden in the oligomeric environment, we carried out a system-environment analysis using the *ProDy* interface [39, 40]. System-environment analysis previously developed for the study of membrane protein in the presence of membrane as environment [41] permits us to view the dynamics of the monomer (system) within the oligomer (environment), and to compare it to the mode spectra available to the monomer in isolation. Our mode-mode correlation analysis shows that the monomers embedded in the restrictive environments of the hexameric or dihexameric complexes still have access to modes of motions exhibited by the monomer in isolation, even if these were not discernible in the soft-mode regime of the oligomers (**Fig. S4A**). Alternatively, one can also examine the correlation between the normal modes of a monomer in isolation and those of the multimers - hexamer or dihexamer. The latter analysis (**Fig. S4B**) clearly shows that the normal modes of the isolated monomer are redistributed and shifted to a higher-frequency regime in the spectrum of modes accessible to the multimers. The multimers have their own low-frequency motions that take precedence.

Overall, **Fig. S4** shows how the monomers in the dihexameric assembly retain the characteristics of the isolated monomer (panels **A** and **B**), but these are not effectively expressed in the soft modes of motions preferentially sampled by the large oligomeric constructs (panel **B**), resulting in the behavior presented in **Fig. 3C–D**. In this respect, it is of interest to see the mean-square fluctuation profiles of residues driven by *all modes* accessible to the intact structures (not only the soft modes), and also to examine the behavior of all monomers in the oligomeric assemblies (not only the first monomer). **Fig. S5** shows the results for the (chains A and B of the) six monomers within the hexamer. Experiments and theory show reasonable agreement, given the coarse-grained representation of the oligomers by the GNM theory, as well as the known effects of non-biological intermolecular contacts on X-ray crystallographic B-factors which preclude an unbiased comparison [42, 43]. The curves show the distinctive behavior of monomers 1-3 from monomers 4-6. Monomers (1,4), (2,5) and (3,6) represent symmetry-related copies of the single same asymmetric dimer within the hexameric assembly.

### GNM analysis of detemir dimers assembled in dihexamers highlights the residues dominating dimer conformational mechanics and stabilization

The fluctuation profile of the dihexamer is more complex. **Fig. S5** displays the behavior of the corresponding 600 residues (100 residues per dimer) clearly showing two types of dimer behavior: dimers 1-3 share the same (*Type 1*) profile and dimers 4-6 the *Type 2*, originating from the different inter-hexamer contacts experienced by the respective hexameric subunits 1 and 2. **Fig. 6, S6** and **S7** present close-up views of the behavior of the respective dimer types (1 and 2) within the dihexamer.

**Figure 6.**
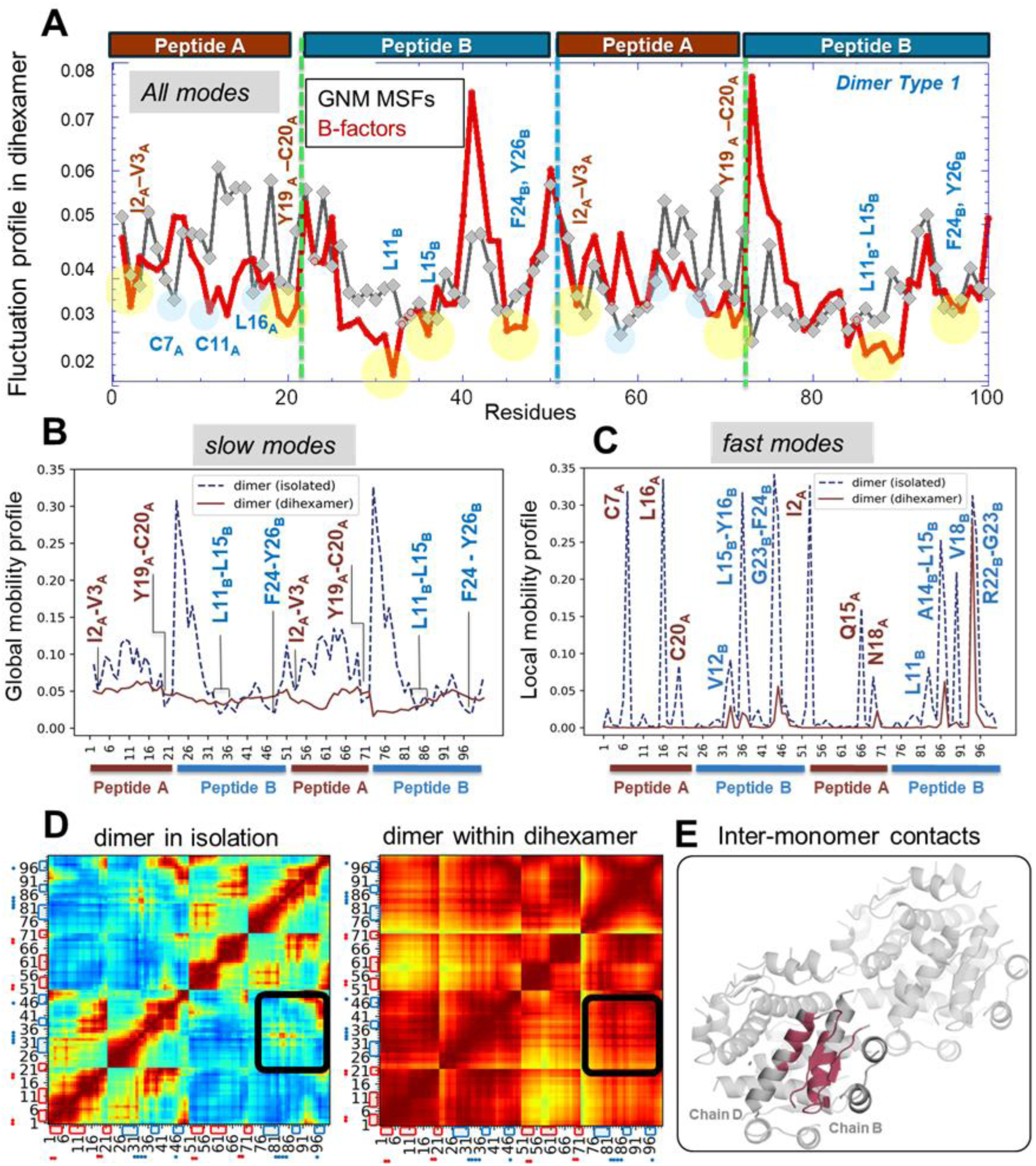
Equilibrium dynamics of the detemir dimer within the dihexameric INSD. The results refer to dimer type 1; those for dimer type 2 are in **Fig S7**. (**A**) Distribution of mean-square fluctuations of residues in dimer of type 1 predicted by GNM (all modes accessible to the dihexamer) and indicated by experimental B-factors. Minima highlighted in *yellow* (and labeled) refer to hinge sites that mediate the collective dynamics, endowed by the global dynamics of the insulin monomer (see **Figs. 3D** and **4**) (**B**) Comparison of the global dynamics of the detemir dimer embedded in the dihexamer with that of the dimer in isolation. **(C)** Comparison of the local dynamics (driven by the fastest GNM modes) of the same two dimers. **(D)** Cross-correlations driven by the slowest seven modes of the dimer in isolation (*left*) and 14 slow modes of the dimer within di-hexamer forms (*right*). *Black squares* show the cross-correlations at the monomer-monomer interface displayed in panel **E**.

The dimer represents the intermediate state between the inactive detemir packaged in the oligomeric construct and its active state accessible upon release from the oligomer. The fluctuation profiles in **Fig. 6** permit us to distinguish the residues that support the conformational mechanics of the dimer (**Fig 6A-B**) as well as those underlying its stability (**Fig 6C**). Minima in the slow modes of the isolated dimer (**Fig 6B**), which persist in all modes accessible to dihexamer (**Fig 6A**), indicate the hinge residues that consistently mediate the conformational mechanics of the INSD dimer. The residues noted above to serve as hinges or anchors for a monomer (I2_A_–V3_A_, Y19_A_–C20_A,_ L11_B_–L15_B_ and F24_B_–Y26_B_; see **Figs. 3D**), highlighted by *yellow spheres* in **Fig 6A**, robustly retain their role in the dimer embedded in the dihexamer. Other residues noted to be constrained in the global modes of monomers (C7_A_, C11_A_, and L16_A_; *light blue spheres*) also lie at the minima of the mean-square fluctuation profiles based on all modes.

Notably, if we solely focus on the global motions of the overall dihexamer (based on its 14 softest modes; see **Fig. S2**), the intrinsic dynamics of the isolated dimer (shown by the *dashed curve* in **Fig 6B**) is hardly observable, as seen in **Fig. 6B**. Likewise, the cross-correlation map generated for the dimer in isolation (**Fig. 6D** *left*) sharply departs from that within the dihexamer (**Fig. 6D** *right*). However, when we consider *all modes* of the dihexamer, the hidden intrinsic dynamic characteristics resurface as seen in **Fig. 6A**. The **Fig. 6E** *(ribbon diagram*) displays the position of the adjacent peptides B of the dimer, which make inter-monomer contacts (highlighted by the *black squares* in the cross-correlation maps). The *left map* shows the opposite-direction movements of the two helices, and consequent high-susceptibility of the two peptides B to dissociate in the isolated dimer. In contrast, they are strongly coupled and move coherently in the dihexameric environment, indicating their lower probability to dissociate, as required for the long-acting property of the INSD dihexamer.

As a further test, we examined the high-frequency end of the mode spectrum (**Fig. 6C** and **Fig. S7C**). Consistent with the results for the monomer (**Fig. 4**), we see among them the disulfide-bridge-forming residues C7_A_, C20_A_ as well as previously noted highly constrained L16_A_. These residues, in accord with their role in the monomer, support the stability of the dimer. Our analysis also invites attention to the dual role of selected hinge residues (minima in slow and all modes) and hot residues (maxima in fast modes); mainly I2_A_, C7_A_, L16_A_, C20_A_, L11_B_, and L15_B_ support both the conformational mechanics and the stability of the dimers within the dihexameric assemblies. Other peaks in the fast modes, which neighbor hinge residues include Q15_A_, N18_A,_ V12_B_, A14_B_, V18_B_, R22_B_-G23_B_-F24_B_ in at least one of the monomers of either dimer type. GNM computations repeated by including additional nodes for the myristoyl groups yielded very similar (correlation coefficient of 0.97) curves, with the major effect being a decrease of the mobility of peptide B terminal residues that make contacts with the myristoylated K29_B_.

The above analysis provided the identity of several residues that help mediate the collective dynamics of the dimeric unit embedded in the dihexamer, as well as those maintaining the stability of detemir in different multimeric states. To make an assessment of the potential involvement of these key residues in the recognition of, or binding to, the insulin receptor, following the release of the dimers and monomers from the oligomeric assemblies, we utilized as reference the cryo-EM structure IR (insulin receptor) complexed with four insulin molecules (PDB ID: 6PXV) [29]. As mentioned in the description of **Fig. 1**, IR harbors two sites (Sites 1 and 2) for insulin binding. **Fig. 7** illustrates the interfacial interactions at the high-affinity site (Site 1). The interactions of peptides A and B are displayed in the respective panels **A** and **B**, obtained by rotating the view by 180°. Notably, the majority of the residues distinguished above by their critical roles in maintaining stability of supporting the global dynamics are critically positioned at interfacial regions.

**Figure 7.**
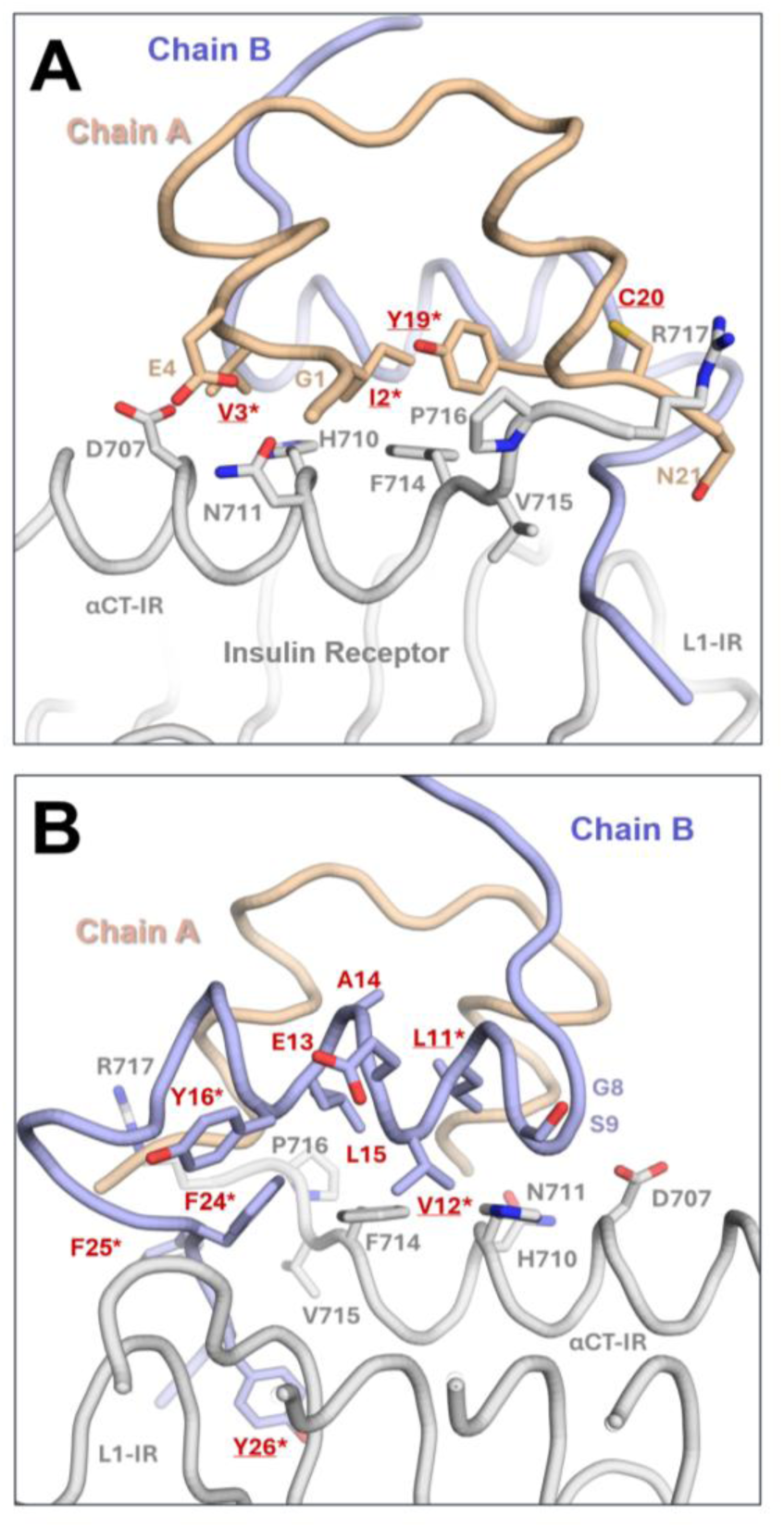
Insulin-insulin receptor interface and involvement of key residues. The insulin-IR interface is shown using cryo-EM structure resolved by Uchikawa et al (PDB: 6PXV) [29]. Panels (**A**) and (**B**) display the respective *front* and *back views*, focusing on the respective interfacial interactions of the peptides A and B of the insulin. Chain A is colored *wheat*; and chain B, *light blue*. The chains α−CT and L1 of IR are colored *gray*. In panel **A**, Chain A residues that interact with the IR (G1_A_, I2_A,_ V3_A_, E4A, Y19A, N21A) are shown in wheat sticks with wheat labels; those identified here to be key to maintaining function are indicated by the *red* labels; I2_A_, V3_A_ and Y19_A_ further overlap with canonical residues interacting with the receptor, as emphasized by an asterisk. The interfacial IR residues (D707, H710, N711, F714, V715, P716, R717) are in *gray sticks* with *gray* labels. In panel **B**, peptide B key residues identified by the GNM to play key roles (L11_B_, V12_B_, E13_B_, A14_B_, L15_B_, Y16_B_, F24_B_, F25_B_, and F26_B_ are labeled *red*. L11_B_, V12_B_, Y16_B_ and F24_B_ -F26_B_ (shown by asterisks) overlap with canonical receptor-binding residues. The IR residues interacting with insulin (D707, H710, N711, F714, V715, P716, R717) are in *gray stick* and labeled *gray*.

## Discussion and Conclusion

The effect of assembly into dimers or higher oligomers, as well as modifications at chain termini, on the structural dynamics of insulin has been a topic of broad interest due to the pharmacological significance of modulating the action of insulin and designing analogs with improved therapeutic effects. While hexameric insulin (human and bovine) and multiple analogs or modified forms have been resolved starting from the late 1980s, by X-ray crystallography and NMR [4, 6–8, 10, 12, 16], the hexameric and dihexameric structures of insulin detemir at ambient temperature are presented here for the first time. Detemir has been observed by cryo-crystallography to preferentially assume a dihexameric (or dodecameric) state, stabilized via B29-myristoyl bridging across adjacent hexamers) [9, 12]. This dihexamer is the canonical form and the most commonly observed quaternary assembly of detemir. The non-canonical form, on the other hand, corresponds to a solitary R₆ hexamer that persists even when the myristoyl-mediated inter-hexamer association is disfavored. The literature documents dihexamer formation *in crystallo* and *in solution* and also supports a temperature- and ligand-dependent equilibrium between dihexamer and hexamer, which, under certain ambient or packing conditions, can shift toward the non-canonical hexamer [44, 45]. Our study differs from previous studies on insulin structures in that (i) it captures insulin detemir at ambient temperature in two oligomeric states, a dihexamer and a non-canonical hexamer, rather than cryogenic detemir alone or non-detemir hexamers; (ii) the structures were determined using a multi-crystal, ambient-temperature strategy that mitigates cryo-induced lattice artifacts; and (iii) they reveal distinct quaternary packing in rhombohedral R3 (short-c hexamer vs. doubled-c di-hexamer), enabling subunit-level comparisons of cooperative dynamics and receptor-facing hinge residues.

Our detailed *in silico* analysis of intra- and inter-molecular dynamics from the isolated monomer to dihexamer forms noted a distinct trend: the dihexamer exhibits a significantly highest collectivity and cooperativity of internal motions compared to the other multimeric hexameric state, and the monomers embedded in the dihexamer are highly stabilized by the constraints exerted by the hexameric environment (**Fig. 3**). While the hexameric environment also restricts the mobility of insulin dimers (ASU), the latter is significantly stronger due to the role of the network of myristoylated groups that exert additional constraints and heterogeneity. While a fully symmetric structure (such as the hexamer) might be more predisposed to undergo cooperative movements, eventually prompting its disassembly, the dihexameric back-to-front packing (**Figs. 1-2**) imparts a non-symmetrical packing between the dimers, not only distinguishing between the monomers forming the three dimers that form the hexameric unit (**Fig. 6**), but between the dimers themselves, as illustrated in **Fig. S6.** Our study suggests that the quaternary organization of the hexamers and overall heterogeneity of the dihexameric assembly provides an environment significantly more stable than those of hexamers.

It is worth noting, however, that despite its high stability manifested by constraints on monomer conformational flexibility, the intrinsic dynamics of the monomers are retained in the oligomeric constructs as evidenced from mode-mode correlations between the isolated monomer and that embedded in the dihexamer (**Fig. S4**). Examination of the complete spectrum of motions, beyond those visible in the window of soft modes, showed the persistence of the intrinsic character of the monomer, which remains ready to be deployed upon release from the oligomeric environment. **Fig. 4** presents the intrinsic dynamics of insulin monomers, essential to their activation and binding to the insulin receptor.

The GNM analysis revealed the identities of several key residues, and their role in maintaining the cooperative dynamics and/or stability of the monomers as well as dimers (ASU) packaged in the dihexamer. In peptide A, our analysis pointed out two regions important to mediating the global conformational changes: I2_A_-V3_A_ and Y19_A_- C20_A_ (**Figs 3**–**5**) consistently distinguished in different oligomerization states. In addition, the disulfide-bridge forming residues C6_A_, C7_A_, C11_A_ (along with C20_A_) as well as L16_A_ were noted to be strongly constrained, also evidenced by their peaks in high frequency modes of monomers and dimers embedded in the hexameric subunits (**Figs. 4C** and **6C**). In peptide B, residues consistently supporting the global mechanics were L11_B_, L15_B_, and F24_B_-F25_B_-Y26_B_ (**Figs. 4A-B, 5,** and **6A-B**); whereas G8_B_, L15_B_ C19_B_, and Y26_B_ were noted as energy localization centers of the monomers (peaks in fast modes, **Fig. 4C**), complemented by their neighbors V12_B_, Y16_B_, V18_B_, R22_B_, and G23_B_ upon dimerization (**Fig. 6C**). Notably, the hinge residues robustly reproduced at multiple scale, L11_B_, L15_B_, and Y26_B_ (**Figs. 3-5**), were pointed out to be closely associated with the ‘primary binding-site’ engagement when binding to insulin receptor [46], consistent with the usual role of hinge-flexing for accommodating or optimizing protein-substrate interactions.

The key role of several residues identified here corroborates several previous experimental or computational studies. The peaks in the fluctuations driven by fast modes, also called kinetically hot residues [35] usually lie in the protein core and tend to be evolutionarily conserved [34, 35]. Notably, V12_B_, Y16_B_, and F25_B_ have been reported to stabilize the dimer, consistent with the protein core/nucleus-forming tendency of the peaks in fast modes. Previous studies reported that G8_B_, L11_B_, V12_B_, L15_B_, F24_B_, and Y26_B_ form a hydrophobic core [46], consistent with the currently observed peaks in fast modes. While the F24_B_ and Y26_B_ side chains substantially contribute to forming this hydrophobic core, they have distinct (additional) functions during the initial detachment process [47, 48]. Specifically, the side chain of F24_B_ is fully buried forming strong hydrophobic interactions with the L15_B_ side chain [47, 49]. In contrast, the aromatic side chain of Y26_B_ is exposed to water and displays increased flexibility due to its interactions with adjacent water molecules [50–52].

Previous applications of elastic network models to other systems have demonstrated that certain key properties of the monomers are retained in multimers, and they may be even amplified in symmetric assemblies [38, 53, 54]. The key residues distinguished by their role in conformational mechanics or stability in the monomeric and dimeric states retain their intrinsic character in the hexamers, while the hexameric architecture and newly acquired collective motions tend to dampen some movements and endow additional stability. For example, the hinge role of F24_B_ discerned here in the dihexameric state can be traced back to the intrinsic properties of the isolated monomer, which is the active form that binds the cognate receptor. Previous simulations pointed to the F24_B_ hinge movement for enabling the transition of the peptide B C-terminal segment from its closed (inactive) to open (active) conformation to bind to the insulin receptor [46, 55, 56]. Likewise, alanine scanning computations of insulin monomer showed that mutations to alanine at L16_A_ and Y19_A_, L11_B_, L15_B_ (distinguished here by their global hinge or anchor properties), and R22_B_ (a highly constrained energy localization site) are critical to maintaining the fold, and lead to misfolding if substituted by alanine [57]. Previous work also demonstrated that V12_B_, Y16_B_, F24_B_, F25_B_, and Y26_B_, are crucial residues for the dimerization of insulin.

Among the other potentially significant residues, R22_B_ stands out as a surface-exposed, positively charged amino acid located at a critical peak, known for facilitating essential electrostatic interactions that influence the stability of the hIns-IR co-complex during the insulin initial detachment process [27]. It is now established that the motif R22_B_-G23_B_-F24_B_-F25_B_-Y26_B_ is pivotal to controlling the interplay with receptors [58–60]. A high degree of cooperativity is observed here between those residues at the monomer-monomer interfaces of the dimers, and R22_B_-G23_B_ are kinetically hot loci while Y26_B_ is global hinge center.

Notably, the FDA has recently approved the first rapid-acting insulin biosimilar (insulin aspart-szjj) [61, 62], and fixed-ratio co-formulations that pair rapid- with long-acting components (e.g., IDegAsp) are increasingly used in severe hyperglycemia [63, 64], broadening clinical strategies that complement the oligomerization-based mechanisms discussed here. Detemir, on the other hand, was recently discontinued directing efforts to an ultralong acting insulin, icodec. Notably, the dihexameric detemir has a shorter acting effect compared to the trimeric icodec. While order of oligomerization can add delay to the detemir clearance, other pharmacokinetics properties such as slow absorption into blood stream, reversible binding to albumin receptor plays a major role in insulin long residence time [65]. Therefore, engineering a slower acting insulin analog does not necessarily require constructing a bulkier molecule but modifying insulin to increase its stability, e.g., by making the substitutions Y14Q_A_, Y16H**_B_**, F25H_B_, in addition to deleting T27_B_ and T30_B_, in icodec [66, 67], and adding C20 fatty diacid side chain for reversibly tighter binding to albumin receptor [68]. Glargine, another type of slow acting insulin, also displays remarkable residence time compared to detemir [69]. Likewise, B29 Lys conjugation with bulky hydrophobic groups, have yielded phenylalanine-conjugated insulin with dramatically improved thermostability against heat, salt, and pH fluctuations [70]. These examples show the ongoing engineering strategies for designing more effective insulin analogs [71], while temperature instability remains a major determinant of insulin treatment failure [72]. The current knowledge of ambient-temperature dihexamer and non-canonical hexamer structures, together with GNM-mapping of critical residues that can be performed for new analogs, may help in engineering tunable next-generation insulins with improved stability, suitable collective dynamics, and optimized therapeutic profiles. In this respect, targeted experimental tests such as site-directed mutagenesis, ligand/ion titrations, solution biophysics are essential to testing and validating the design strategies inferred from computational studies.

## Methods

### Sample preparation and crystallization

Commercially available insulin detemir (Levemir, Novo Nordisk A/S, Bagsvaerd, Denmark) was crystallized using a sitting-drop micro batch vapor diffusion screening technique under oil using 72-well Terasaki crystallization plates [73]. Briefly, the detemir solution was mixed with (1:1 ratio, v/v) ∼3500 commercially available sparse matrix crystallization screening conditions at ambient temperature (293 K) for crystallization. Each well, containing 0.83 µL protein and crystallization condition, was sealed with 16.6 µL of paraffin oil (Cat#ZS.100510.5000, ZAG Kimya, Türkiye) and stored at room temperature until crystal harvesting. Terasaki plates were checked under a compound light microscope for crystal formation and growth. Large crystals were obtained within 48 hours. The best crystals were grown in a buffer containing 0.2 M sodium acetate trihydrate and 0.1 M TRIS hydrochloride pH 9.0.

### Sample delivery and XtalCheck-S setup for data collection

Rigaku’s XtaLAB Synergy Flow XRD system controlled by CrysAlisPro 1.171.42.59a software (Rigaku Oxford Diffraction, 2022) was used for data collection as described in Gul et al. (2023) [25]. The airflow temperature of Oxford Cryosystems’s Cryostream 800 Plus was adjusted to 300 K (26.85 °C) and kept constant for data collection at ambient temperature. Instead of the intelligent goniometer head (IGH), the 72-well Terasaki plate was placed on the modified adapter of the *XtalCheck-S* plate reader attachment mounted on the goniometer omega stage. Two dozen crystals were used for initial screening to rank diffraction quality. Omega and theta angles and then X-, Y-, and Z-coordinates were adjusted to center crystals at the eccentric height of the X-ray focusing region. After centering, diffraction data were collected for each crystal. Well-diffracting crystals were selected for further use in data collection, and the exposure time was optimized to minimize radiation damage.

The best diffracting crystals were grown in a buffer containing 0.09 M HEPES–NaOH, pH 7.5, 1.26 M sodium citrate tribasic dihydrate, 10% v/v glycerol (Crystal Screen Cryo (Cat#HR2-122)). A total of 24 were screened for 9LVX (20 crystals for 9LVY), of which 15 crystals showing the highest diffraction quality (**Table 1, Table S1**) and consistent unit-cell parameters (**Table S2**) for 9LVX and 9LVY were selected for merging, while the remaining crystals were excluded. During data collection, XtalCheck-S was set to oscillate as much as the detector distance would allow to maximize crystal exposure oscillation angles. Diffraction data were collected (∼21° total rotation per crystal, corresponding to 21 frames) for 1 min and 45 s (5 secs/frame) for each run from all individual crystals. Only datasets with consistent diffraction quality and indexing solutions were included in the final merging step, while datasets showing inconsistencies or lower quality were excluded. A total of 15 crystals were used for the final merged dataset for both dataset (**Fig. S3**). The detector distance was set to 100.00 mm, the scan width was set to 1.00-degree oscillation, and the exposure time was set to 5.00 s per image.

### Data processing

Once the plate screening parameters were optimized for all crystals, 21 degrees of data collection were performed for each prescreened/selected crystal. All crystals were queued in CrysAlisPro for complete data collection. An optimal unit cell was chosen, and peak finding and masking were performed for the data collected. A batch script was generated with the xx *proffitbatch* command for cumulative data collection. The batch data reduction was run on CrysAlisPro Suite using the script command. Data reduction produced a file that contains all integrated, unmerged, and unscaled data (*.rrpprof) for each dataset. For merging all datasets as a reflection data (*.mtz) file, the proffit merge process from the Data Reduction section on the main window of CrysAlisPro was used. Reduced datasets (*.rrpprof files) were merged again using proffit merge as described. All data was refinalized, merged, and scaled with aimless and pointless implementation in CCP4. Finally, the processed data was exported as *.mtz format.

### Structure determination

The crystal structures of INSD structures were determined at ambient temperature in space group R3:H by using the automated molecular replacement program *PHASER* implemented in the *PHENIX* software package [74]. A previously published X-ray crystal structure was used as an initial search model (PDB ID: 8HGZ) [9]. 8HGZ structural coordinates were used for the initial rigid-body refinement within PHENIX. After simulated-annealing refinement, individual coordinates and Translation/Libration/Screw (TLS) parameters were refined. Additionally, composite omit map refinement implemented in PHENIX was performed to identify potential positions of altered side chains and water molecules. The final model was checked and rebuilt in COOT version 0.8.9.2 [75], while positions with a strong difference density were retained. Water molecules located outside of significant electron density were manually removed. All X-ray crystal structure figures were generated with *PyMOL* version 2.3 [76] and *COOT*.

### Gaussian Network Model analysis of INSD dynamics

GNM analysis was performed using the *ProDy* package [39] for two newly determined INSD structures at dihexameric and hexameric states, as well as their monomeric and dimeric components, and the isolated monomers and dimers. The approach is based on the construction and eigenvalue decomposition of the Kirchhoff matrix **Γ** (N x N, N being the total number of residues) that describes the connectivity of the network. The cutoff distance for long-range interaction was set at 8.0 Å. GNM yields N-1 nonzero normal modes the frequencies and shapes of which are defined by the respective eigenvalues and eigenvectors of **Γ**. The cross-correlations between the equilibrium motions of the residues are obtained from the pseudoinverse of **Γ,** organized in a matrix, **C**, the diagonal elements of which are the mean-square fluctuations of residues, and off-diagonal terms are the cross-correlations. Normalization of the cross-correlations by mean-square fluctuations of the residues yields the orientational cross-correlations. **C** may be written as a summation over the contributions of the individual modes of motions obtained from the eigenvalue decomposition of **Γ** (or **C**). The heat maps presented in this study (**Figs. 3B-C, 6D, S3,** and **S4C**) represent the normalized cross-correlations, also called correlation cosines between pairs of residue motions, contributed by the softest modes.

The inverse eigenvalues of **Γ** serve as the weight of individual modes; the relative contributions of the individual modes are obtained by evaluating the fractional contribution of each mode; and the cumulative contribution/variance for subsets of modes (**Fig. S2**) by their sum. The softest modes at the lowest frequency end of the mode spectrum were selected to contribute a minimal variance of 0.40, and the fastest modes were represented by the 10 highest frequency modes at the other end of the spectrum. System-environment analyses on monomers embedded within the oligomers were performed by considering the system Kirchhoff matrix Γₛₛ, computed by [Γ_ss_ − Γ_se_ Γ_ee_^⁻¹^ Γ_se_^T^] where the subscripts indicate the submatrices of Γ corresponding to intra- and inter-interactions of the system (s) and environment (e). Mode-mode correlations (**Fig. S2)** are evaluated by computing the correlation cosines between the modes (represented by N-dimensional eigenvectors) accessible to the isolated structure (e.g. monomer) and to the same structure embedded in an environment. Mrystol moieties have been included by extending the set of nodes used to build the network through atom selections that retain these groups and by supplying custom coordinates for additional nodes. The resulting contacts are then determined from the input structure’s geometry and topology.

## Ethics approval and consent to participate

Not applicable.

## Availability of data and materials

The coordinates and structure factors have been deposited in the Protein Data Bank under the accession codes 9LVX (dihexamer form of detemir) and 9LVY (non-canonical hexamer form of detemir).

## Authors’ contributions

E.A., T.H., and I.B. designed the study. E.A. prepared the samples. E.A. performed the sample delivery, data collection, data processing, and structure determination under the supervision of H.D; and E.A. and H.N. performed the computations under the supervision of T.H., and I.B. The manuscript, originally written by E.A., H.N., T.H., H.D., and I.B, was finalized by H.N. and I.B.

## Acknowledgment

The authors gratefully acknowledge the use of the services and facilities of the University of Health Science-Validebag DETAUM (Experimental Medicine Research and Application Center). They also thank Soichi Wakatsuki for numerous discussions regarding the interpretation of the 9LVX structure in terms of its biological rather than asymmetric unit, considering the symmetry-averaged nature of the special R3:H space group. IB gratefully acknowledges support from the National Institutes of Health grants R01 GM139297 and R01 DK116780.

## Supplementary Material

### Supplementary Figures

**Figure S1.**
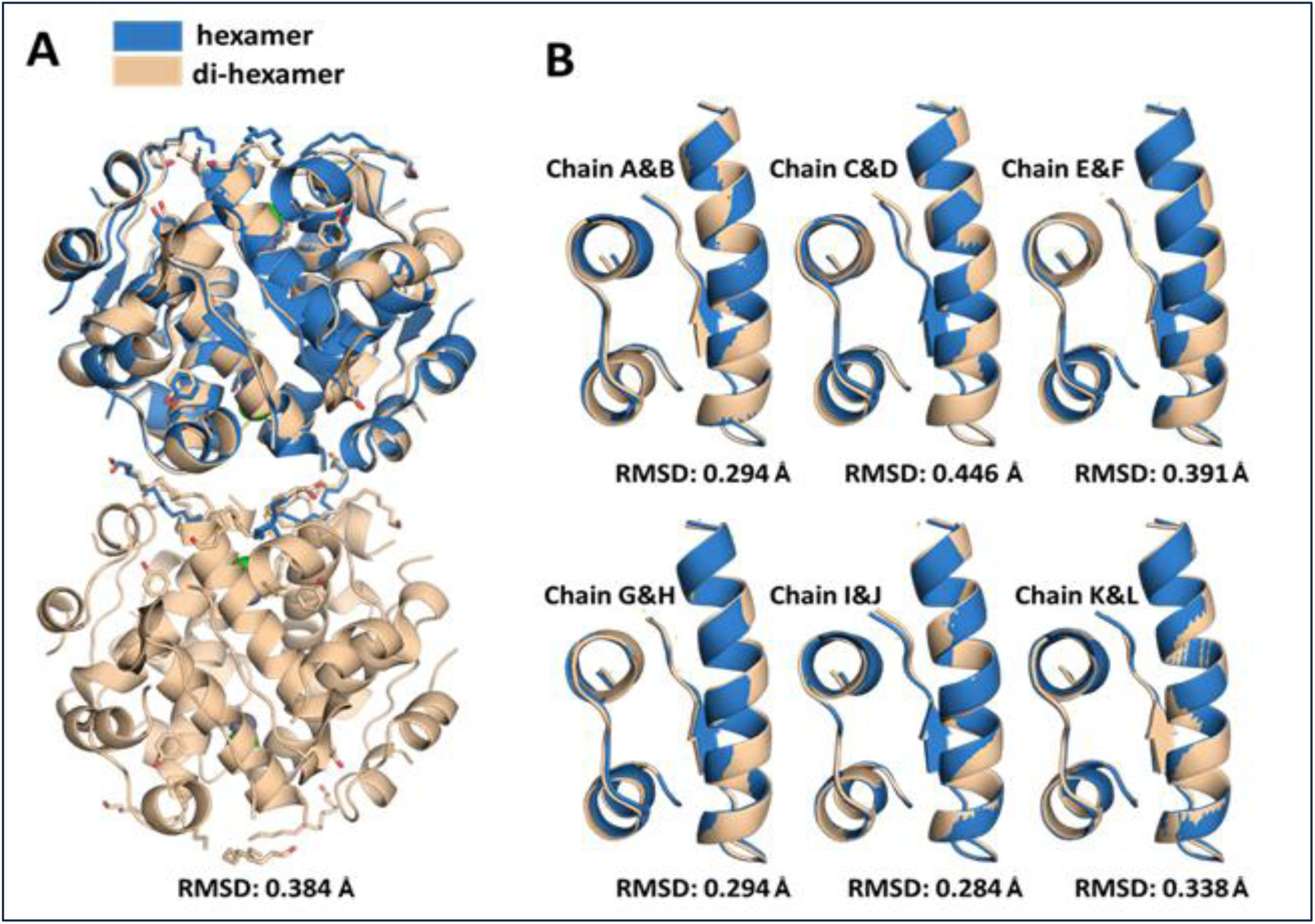
Alignment of the rhombohedral crystal structure of dihexamer INSD against the hexamer form, and the role of myristoyl chains at dihexamer interface. **(A)** Superposition of dihexamer and non-canonical hexamer INSD. **(B)** Superposition of the individual monomers within the dihexamer and hexamer INSD.

**Figure S2.**
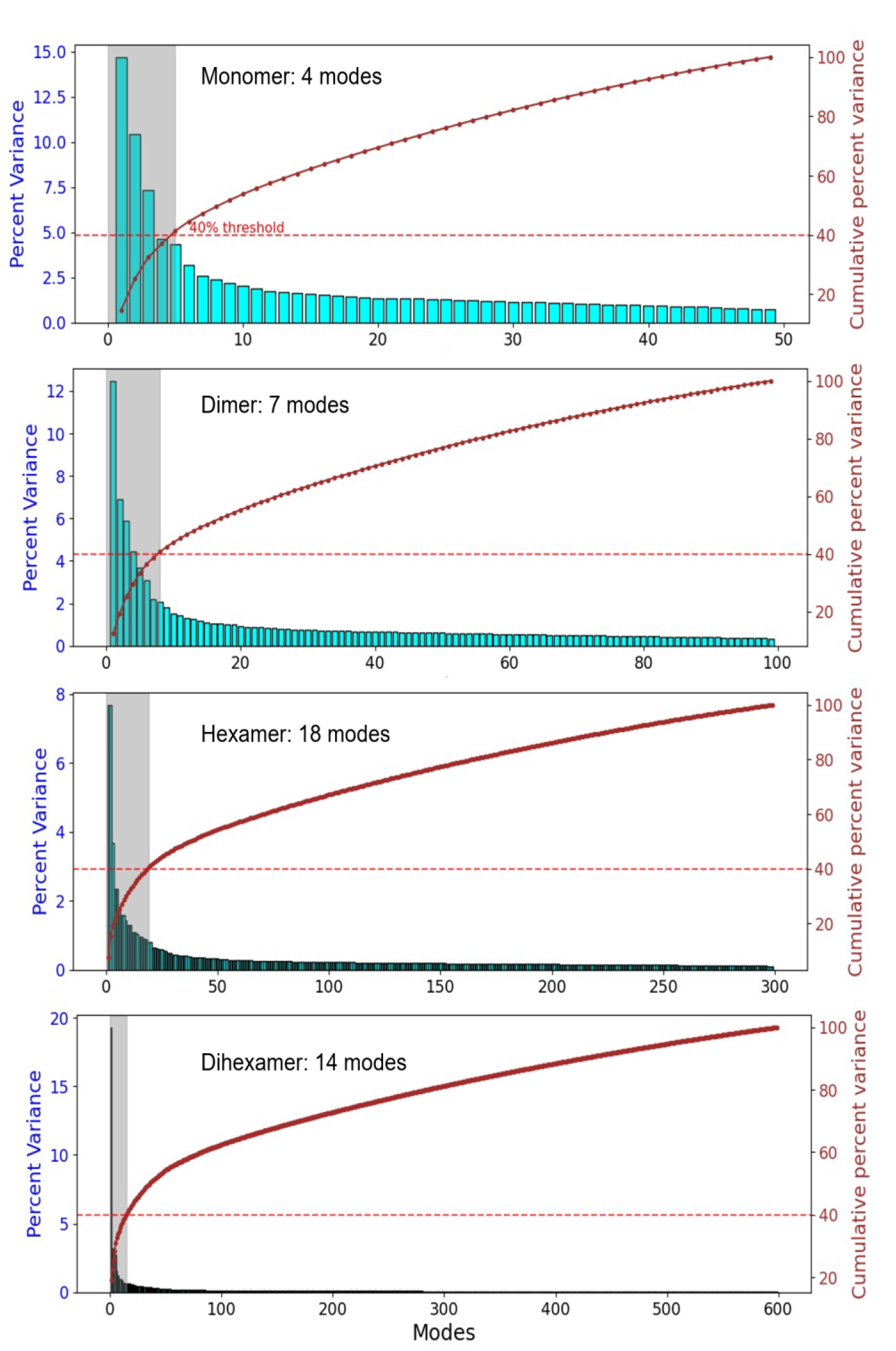
Percent contribution of GNM normal modes to the complete spectrum of motions in different oligomers. Percent variance of each mode is shown by *cyan bar*s (*left* ordinate); whereas the cumulative variance is shown by the *red curve* with *brown dots (right* ordinate). The *shaded grey box* highlights the modes contributing to the soft modes in each case, the cumulative contribution of which makes up > 40% of the total variance, as indicated by the dashed horizontal line which refers to the right ordinate. We note that in the dihexamer, a single mode (the slowest mode) makes a major contribution (of ∼ 19%). This mode refers to the relative movements of the two hexamers, each moving as a rigid block.

**Figure S3.**
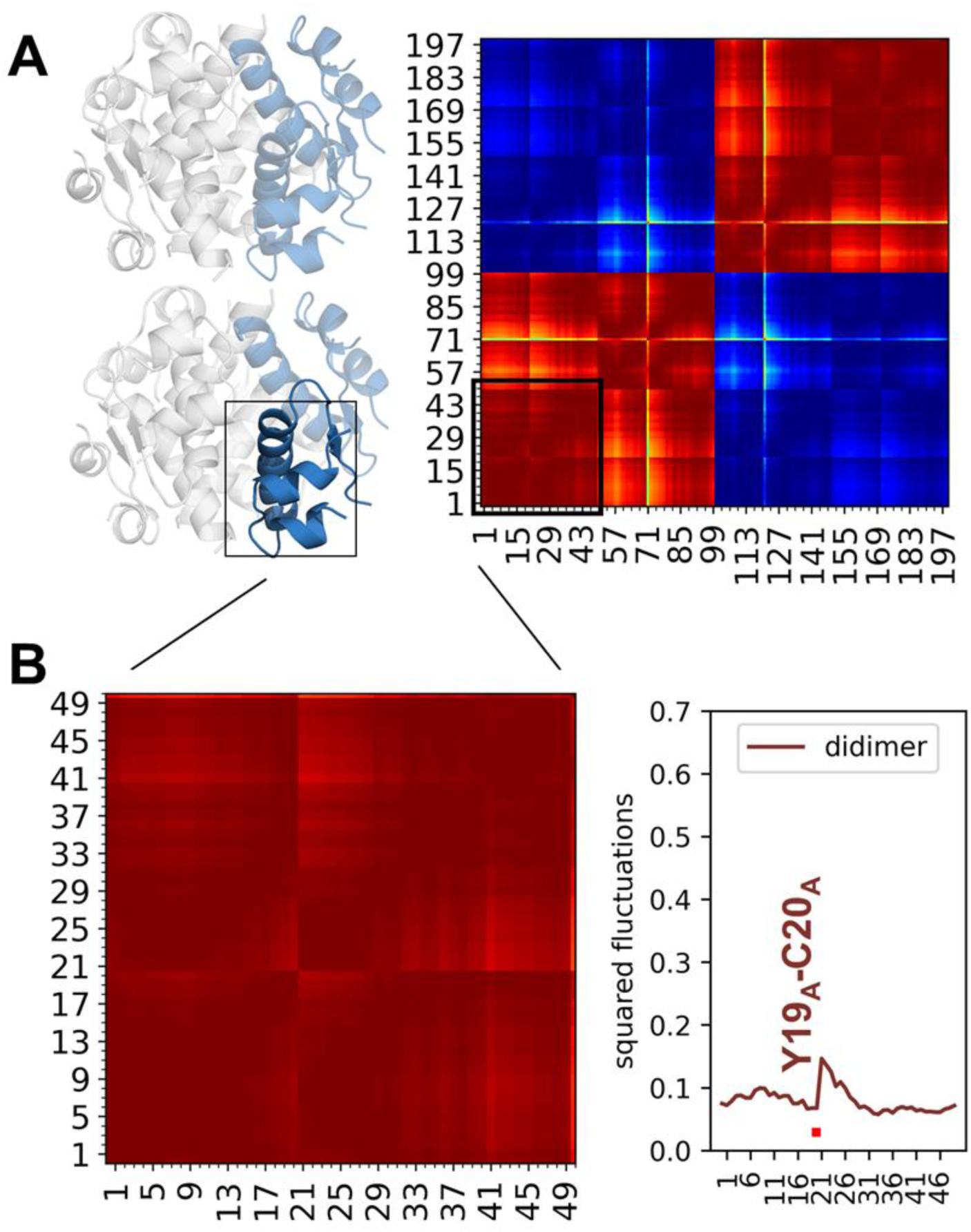
Intrinsic dynamics of tetramers composed of two dimers (also called di-dimer) lying each in a different hexamer. Panel **A** shows the di-dimer (*blue*) and a monomeric subunit (*dark blue*) within the di-dimer. The corresponding cross-correlation map based on the slowest GNM modes (amounting to 40% cumulative variance) is shown on the *right*. The map is color-coded by the correlation cosines of residue pair motions, ranging from −1 (dark blue, fully anticorrelated; coupled movements in opposite directions) to +1 (fully correlated; coupled movements in the same direction). The map clearly shows the anticorrelated movements of the two pairs of dimers, while the monomers within each dimer are strongly correlated as enlarged in panel **B**. Panel **C** shows the mobility profiles of the highlighted monomer, where the y-axis represents the mean-square fluctuations normalized for the structure to which the monomer belongs, and the x-axis represents the residue numbers.

**Figure S4.**
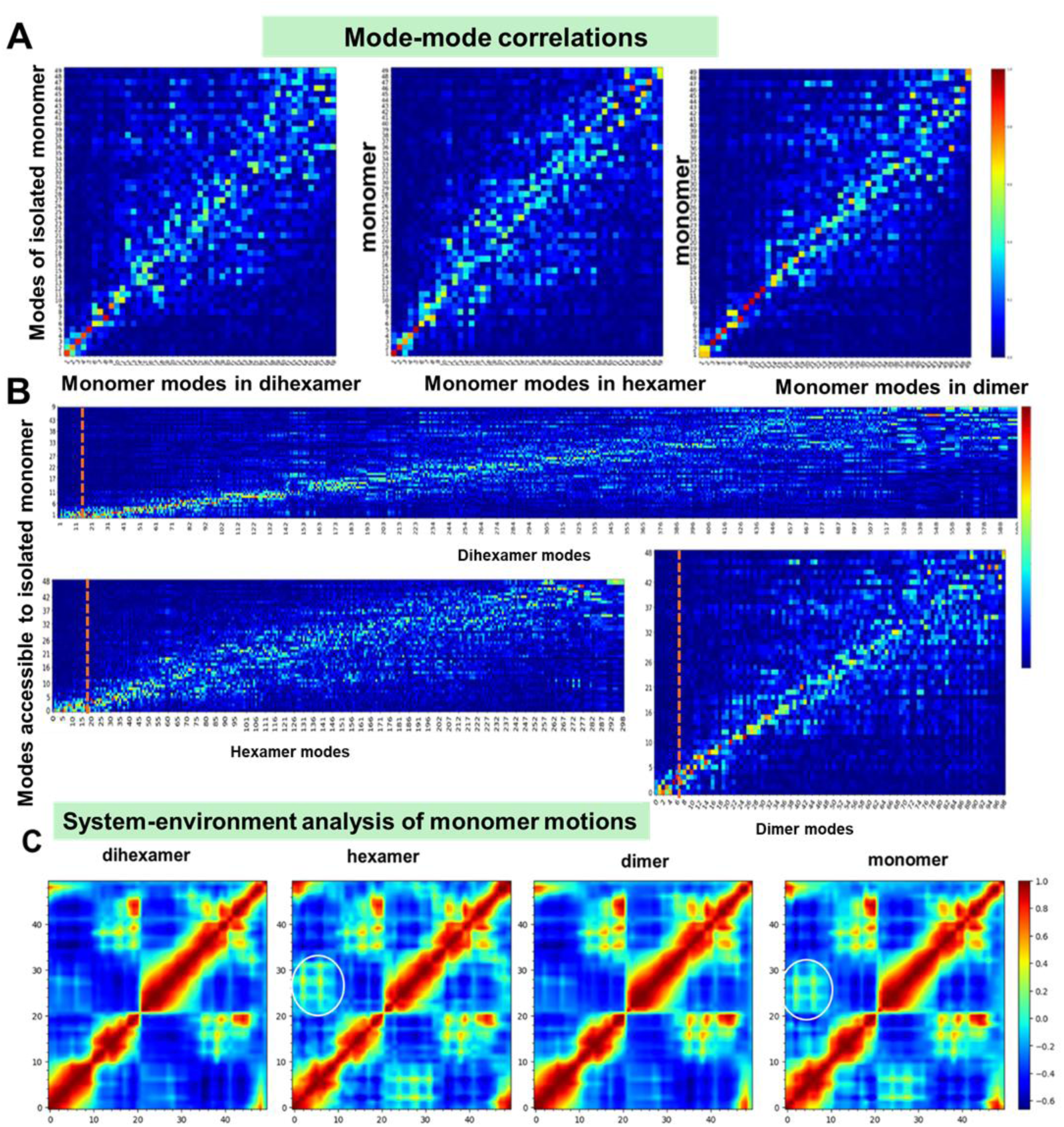
Results from system-environment analysis of the dynamics of INSD monomers when taking part in different multimers. **(A)** Mode-mode correlations between the first monomer of the multimers (x-axis) and the isolated monomers (y-axis). Mode-mode correlations are evaluated by calculating the absolute value of the correlation cosines between the modes accessible to the isolated monomer, and those accessible to the first monomer of multimer.The modes are described by the eigenvectors of the GNM Kirchhoff (or connectivity) matrix evaluated using *ProDy* tools for both the isolated monomer, and for the first monomer of the multimers. The latter is treated as the ‘system’ whose dynamics is mediated by the ‘environment’ created by the other monomers of the multimer. The mode-mode correlations vary from *dark blue* (no correlation) to *dark* red (fully correlated). (**B**) Comparison of the mode spectrum of the isolated monomer (y-axis) to all modes accessible to the dihexamer, hexamer and dimer. The *orange vertical bar* indicates the cutoff for soft modes of the oligomers adopted in **Figure 3** (see also **Figure S1**). The modes of the monomers are moved to higher indices (higher frequency regime) in oligomers, highlighting the conservation of monomers’ modes of motion but their lower contribution to the oligomer dynamics as new modes of motion unique to oligomers emerge. **(C)** Inter-residue cross-correlations based on the first 10 modes accessible to the monomers (systems) embedded in the restrictive environment, after removal of global modes of the environment, revealing the conservation of the monomer intrinsic dynamics. Circled regions highlight the small changes across the different oligomeric states.

**Figure S5.**
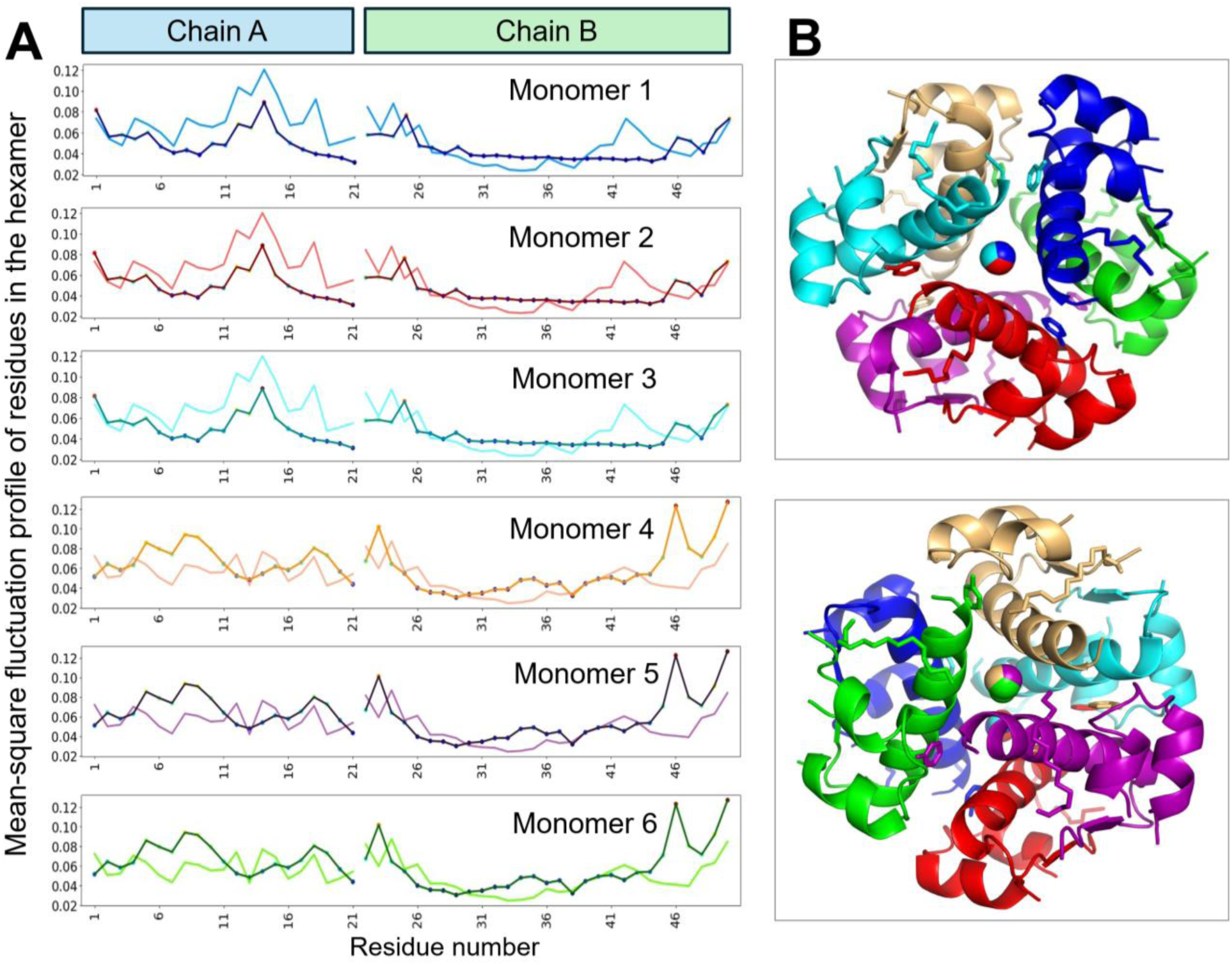
Equilibrium fluctuation profiles of detemir monomers in the hexamer, based on all modes of motions accessible to the hexamer. **(A)** Two normalized curves are displayed for each monomer: (i) the distributions of MSFs evaluated for each detemir monomer (including its myristoyl terminal group) using the complete spectrum of GNM modes predicted for the hexamer (*lighter curve*), and (ii) the X-ray crystallographic B factors (*darker curve, with dots*). The curves are colored by the corresponding monomers in the ribbon diagrams shown in panel **B**. Monomers 1-3 (B *top diagram*; monomers in the front) share the same profile, as well as monomers 4-6 (panel **B**, the *bottom diagram front monomers*).

**Figure S6.**
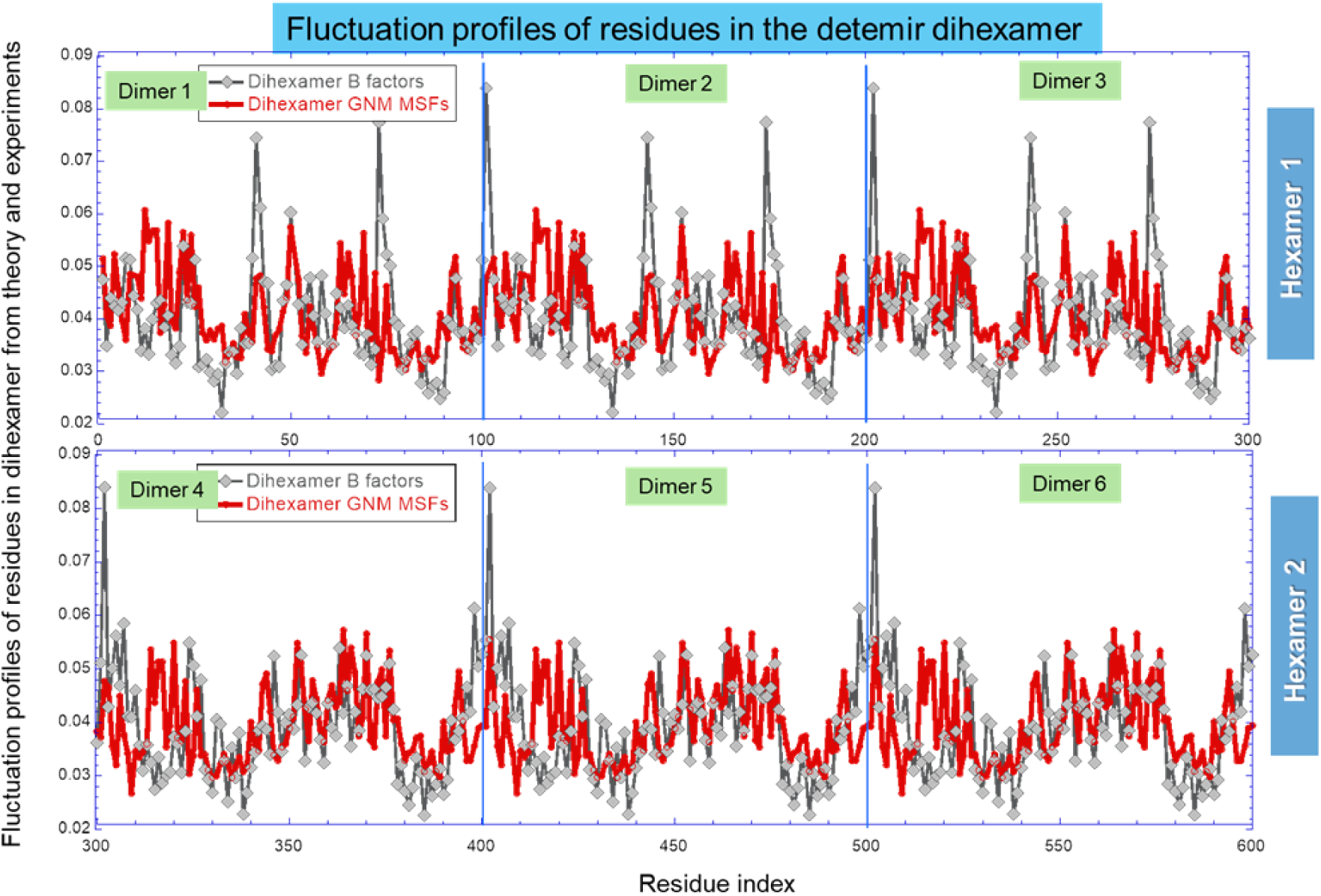
Fluctuation profile of residues in the INSD dihexamer. The top and bottom panels display the behavior of two hexamers (respective hexamer 1 and 2, labeled on the *right*), each comprising three dimers of 100 residues (delimited by *vertical lines*, and *labeled*). The two hexamers obey different dynamics due to their asymmetric interfacial contacts originating from the back-to-front stacking of the hexamers. The mean-square fluctuation profile predicted by the GNM (all modes; *red curve*) and the B-factor measured in the crystal (*gray curve* with *diamond symbols*) are displayed. Both curves are normalized. A closeup view of the plot for the first dimer of hexamer 1 is presented in **Fig. 6A**. GNM computations repeated by including additional nodes for the myristoyl groups yielded very similar (correlation coefficient of 0.97) curves the major effect being a decrease of the mobility of peptide B terminal residues that make contacts with the myristoylated K29_B_. Residues are numbered from 1 to 600 along the x-axis, referring to the 6 dimers of 100 residues each. Each dimer is composed of 2 distinct monomers, and each monomer, of two peptides A (21 residues) and B (29 residues)

**Figure S7.**
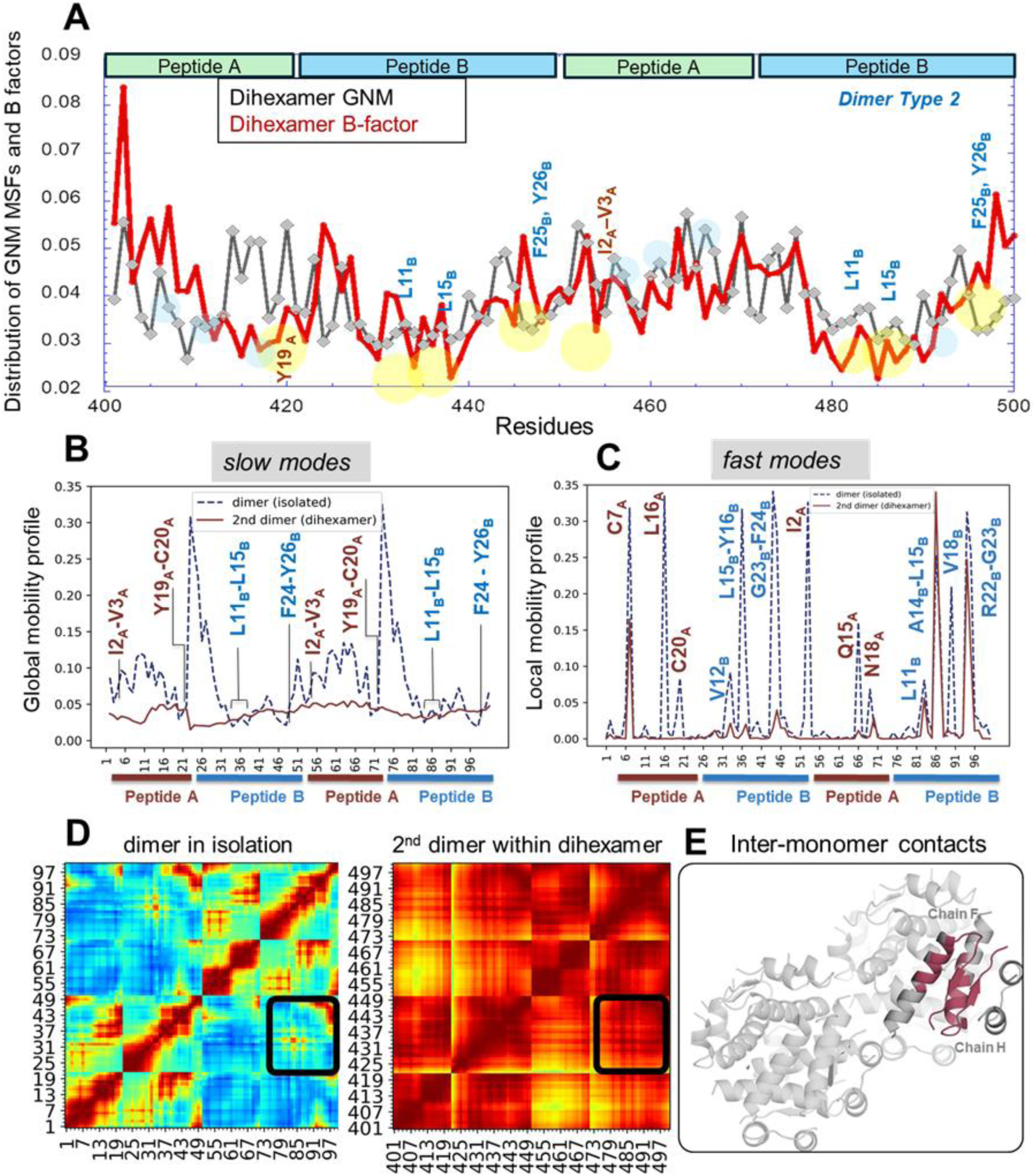
Equilibrium dynamics of the detemir dimer of type 2 within the dihexameric INSD. The figure is the counterpart of main text **Fig. 6**, repeated for dimer of type 2 embedded in the dihexamer. (**A**) Distribution of mean-square fluctuations of residues predicted by GNM (all modes accessible to the dihexamer) and indicated by experimental B-factors. Minima highlighted in *yellow* (and labeled) refer to hinge sites that mediate the collective dynamics, endowed by the global dynamics of the insulin monomer. (**B-C**) Comparison of the global (**B**) and local (**C**) dynamics of this detemir dimer (embedded in the dihexameric state) with those of the dimer in isolation. **(D)** Cross-correlations driven by the slowest seven modes of the dimer in isolation (*left*) and 14 slow modes of the dimer within di-hexamer forms (*right*). *Black squares* show the cross-correlations at the monomer-monomer interface displayed in panel **E**.

